# Nascent alt-protein chemoproteomics reveals a repressor of ribosome biogenesis

**DOI:** 10.1101/2021.06.29.450363

**Authors:** Xiongwen Cao, Alexandra Khitun, Cecelia M. Harold, Carson J. Bryant, Shu-Jian Zheng, Susan J. Baserga, Sarah A. Slavoff

**Affiliations:** Department of Chemistry, Yale University, New Haven, Connecticut 06520, United States; Institute of Biomolecular Design and Discovery, Yale University, West Haven, Connecticut 06516, United States; Department of Genetics, Yale University School of Medicine, New Haven, Connecticut 06520, United States; Department of Molecular Biophysics and Biochemistry, Yale University, New Haven, Connecticut 06520, United States; Department of Therapeutic Radiology, Yale University School of Medicine, New Haven, Connecticut 06520, United States

## Abstract

Many unannotated microproteins and alternative proteins (alt-proteins) have recently been found to be co-encoded with canonical proteins, but few of their functions are known. Motivated by the hypothesis that alt-proteins undergoing active or stress-induced synthesis could play important cellular roles, here, we developed a chemoproteomic pipeline to identify nascent alt-proteins in human cells. We identified 22 actively translated unannotated alt-proteins, one of which is upregulated after DNA damage stress. We further defined MINAS-60 (*MI*croprotein that *N*egatively regulates *AS*sembly of the pre-*60*S ribosomal subunit), a nucleolar localized alt-protein co-encoded with human RBM10.Depletion of MINAS-60 increases the amount of the mature 60S ribosomal subunit, consequently upregulating global protein synthesis and cell proliferation by repressing late-stage pre-60S assembly and export of the 60S ribosome subunit to the cytoplasm. Together, these results implicate MINAS-60 as a repressor of ribosome biogenesis, and demonstrate that chemoproteomics can enable generation of functional hypotheses for uncharacterized alt-proteins.

## Introduction

Expression of thousands of previously unannotated small open reading frames (smORFs, typically defined as ORFs comprising fewer than 100 codons and, in some studies, up to 150 codons^1^) has recently been revealed in mammalian cells^2^. These smORFs are found in long non-coding RNAs, 5’ and 3’ untranslated regions (UTRs) of mRNAs, and frame-shifted ORFs overlapping protein coding sequences (CDS), the latter of which are termed alternative ORFs or “alt-ORFs”^3^. A rapidly increasing number of smORF-encoded proteins (SEPs, also known as micropeptides and microproteins) and alt-ORF-encoded proteins (alt-proteins)^2^ have been shown to play important roles in vertebrate biology. For example, the human SEPs BRAWNIN and MOCCI function in oxidative phosphorylation^4,5^, and MP31 is a tumor suppressor that regulates lactate metabolism in glioblastoma^6^.

Fewer alt-proteins have been well-defined, including alt-FUS, which cooperates with FUS in formation of cytotoxic aggregates^7^, and alt-RPL36, which regulates the PI3K-AKT-mTOR pathway^8^. Recently, genome-scale CRISPR screens revealed that hundreds of smORFs regulate human cell growth and survival^9,10^. These studies demonstrate that defining the function of bioactive SEPs and alt-proteins represents a major opportunity to gain insights into biological complexity.

Currently, there is proteomic and ribosome profiling evidence for tens of thousands of human smORFs and alt-ORFs^3,11^, but the overwhelming majority of smORFs and alt-ORFs remain uncharacterized, in part because their short lengths and, in some cases, limited conservation render homology-based annotation challenging^12^. Furthermore, most alt-ORFs remain unstudied because it is challenging to separate their functions from the canonical protein CDS in which they are nested. Because strategies to identify alt-proteins that participate in biological processes are currently in the developmental stages, it remains unclear whether alt-proteins are broadly functional. We hypothesize that alt-proteins with properties (e.g., chemical reactivity, regulated expression) similar to those of canonical proteins are likely to play important cellular roles, and that chemoproteomic strategies targeted to those properties can be leveraged to identify functional alt-proteins. Providing precedent, a chemoproteomic profiling study of microproteins containing reactive cysteine residues, which is a feature of the active site of many enzymes, revealed 16 nucleophilic microproteins, though their cellular roles were not explored in that study^13^. In this work, we test the hypothesis that alt-proteins undergoing active or stress-induced synthesis are likely to be functional, and we develop a chemoproteomic approach to identify them.

Using this method, we identify an alt-ORF that overlaps the human RBM10 CDS and encodes a repressor of ribosome large subunit (LSU) biogenesis. Ribosome biogenesis is a highly spatially and temporally regulated cellular process essential for growth and development^14-17^. In humans, ribosome biogenesis starts in the nucleolus with the transcription of a 47S precursor rRNA (pre-rRNA) by RNA polymerase I (RNAPI). The 47S pre-rRNA is chemically modified and complexed with ribosome assembly factors and ribosomal proteins to form the 90S pre-ribosomal particle. Endonucleolytic cleavage of the 47S pre-rRNA subsequently generates the pre-40S and pre-60S particles, which undergo individual maturation and quality-control steps prior to export to the cytoplasm.

Pre-60S ribosomal particles have recently been probed by cryo-electron microscopy (cryo-EM)^18,19^, revealing a sequential maturation process involving quality control checkpoints for 5S ribonucleoprotein particle incorporation and rotation^20^, active site formation, and removal of internal transcribed spacer 2 (ITS2) prior to export from the nucleus to the cytoplasm. In the cytoplasm, the final steps of maturation occur to produce large 60S and small 40S subunits, which associate to form translation-competent ribosomes. Dysregulated ribosome biogenesis has been linked to numerous human disorders, including cancer^21^, Alzheimer’s disease^22^, and congenital disorders termed ribosomopathies^23,24^.

In this study, we developed a chemoproteomic pipeline to identify nascent alt-proteins, which modifies a powerful previously reported strategy for metabolic unnatural amino acid incorporation to be amenable to microprotein enrichment and profiling, and identified 22 unannotated alt-proteins in a human cell line. We confirmed the translation of six selected alt-proteins, and functionally defined one alt-ORF nested in the human RBM10 CDS. We name the encoded alt-protein MINAS-60, or *MI*croprotein that *N*egatively regulates *AS*sembly of the pre-*60*S ribosomal subunit. We show that MINAS-60 localizes to the nucleolus, associates with LSU assembly factors GTPBP4 and MRTO4, and co-fractionates with pre-60S complexes in nuclear extracts. Finally, we engineer MINAS-60 knockdown and rescue cell lines to demonstrate that loss of MINAS-60 increases cytoplasmic 60S ribosome subunit levels, global protein synthesis, and cell proliferation. This is independent of the function of the canonical RBM10 protein, which has been previously shown to inhibit cell proliferation through its role in alternative pre-mRNA splicing^25-29^. These results implicate MINAS-60 as a rare negative regulator of late nuclear steps in LSU biogenesis, and demonstrate that chemoproteomic profiling can prioritize alt-ORFs for functional study.

## Results

### Chemoproteomic profiling of nascent alt-proteins

Motivated by the hypothesis that alt-proteins undergoing active synthesis or stress-induced synthesis could play important cellular roles, we set out to develop a chemoproteomic approach to identify newly translated alt-proteins (Figure 1a). We leveraged bio-orthogonal non-canonical amino acid tagging (BONCAT), an *in vivo* labeling strategy to identify nascent proteins^30^. BONCAT utilizes the methionine analogue azidohomoalanine (AHA), which can be metabolically incorporated into all newly synthesized proteins by the endogenous protein translation machinery. Previous BONCAT workflows included procedures, such as column-based biotin removal, that were likely to eliminate small proteins. We modified the protocol to capture these small proteins, including microproteins and alt-proteins. Labeled (and unlabeled) small proteins were selectively enriched from whole proteome extracts with a C8 column following a previously reported strategy^31^ (Figure 1b), and AHA-containing small proteins were captured with click chemistry directly on dibenzocyclooctyne (DBCO) magnetic beads. On-bead digest was followed with our previously reported liquid chromatography/tandem mass spectrometry-based method for identification of unannotated microproteins and alt-proteins^32^. Using this strategy, we profiled translation in HEK 293T cells under basal growth conditions (DMSO treatment), oxidative stress (sodium arsenite treatment), DNA damage stress (etoposide treatment) and unfolded protein response stress (DTT treatment). In total, we identified 22 unannotated alt-proteins (Figure 1c, Supplementary Data 1), nine of which were specifically detected under stress conditions, including alt-CNPY2, which was specifically detected after etoposide treatment (Figure 1d, Supplementary Data 1).

**Figure 1.**
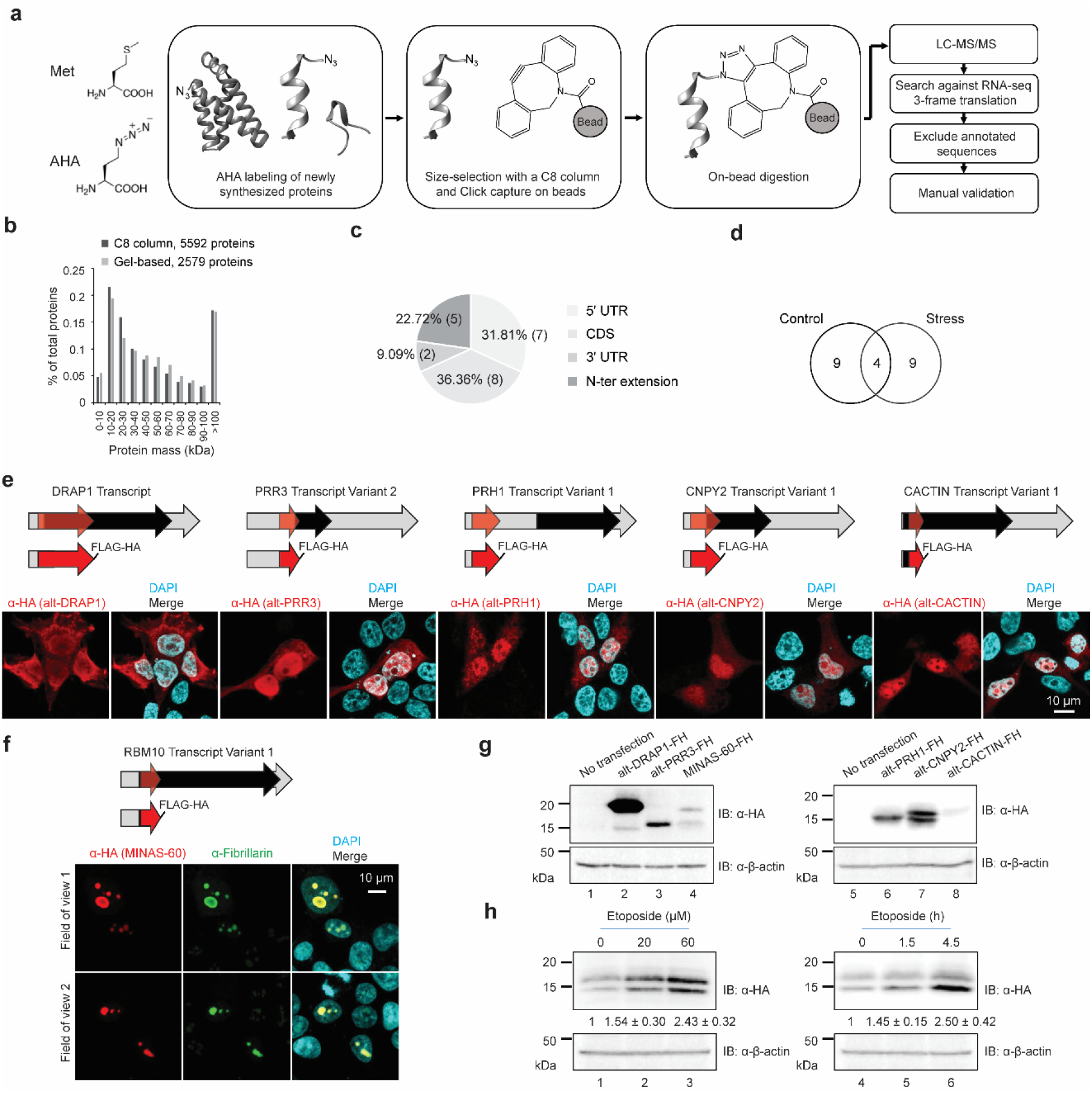
BONCAT-based chemoproteomic identification of newly synthesized alt-proteins. **a** Schematic workflow for BONCAT-based chemoproteomic analysis of nascent alt-proteins. **b** Size distribution of canonical, annotated proteins detected using C8 column-based selection (black) and our previously reported^60^ gel-based selection (gray). **c** Distribution of locations of identified alt-proteins relative to the annotated coding sequence (CDS). **d** Venn diagram of alt-proteins identified under control or stress conditions. **e, f** Top: a schematic representation of human *DRAP1* transcript, *PRR3* transcript variant 2, *PRH1* transcript variant 1, *CNPY2* transcript variant 1, *CACTIN* transcript variant 1, or *RBM10* transcript variant 1 (**c**); light gray arrow, 5’ and 3’ untranslated regions (UTR); red, alternative open reading frame (alt-ORF) coding sequence; black, annotated coding sequence. Middle: a schematic representation of the expression constructs containing the complete 5’UTR and the alt-ORF of the transcript indicated above, with a dual FLAG and HA tag appended to the C-terminus of the alt-protein. Bottom: HEK 293T cells transfected with the expression construct indicated (middle) were immunostained with anti-HA (red), DAPI (cyan), and anti-fibrillarin (green, **c**). Scale bar, 10 µm. Data are representative of three biological replicates. **g** HEK 293T cells transfected with the expression construct (middle) were immunoblotted (IB) with antibodies indicated to the right, with untransfected (no transfection) HEK 293T cells as a control. Data are representative of three biological replicates. **h** HEK 293T cells transfected with the alt-CNPY2 expression construct were treated with increasing amounts of etoposide or vehicle for 2 h (left), or with 60 µM etoposide for different times or vehicle (right), followed by western blotting with antibodies indicated to the right. Quantitative analysis (*N* = 3) of the western blot signal of alt-CNPY2-FLAG-HA are indicated at the bottom.

To confirm the translation and examine subcellular localization of six selected alt-proteins (Supplementary Figures 1a-f, Supplementary Table 1), the cDNA sequence comprising the 5’UTR of the encoding transcript through the stop codon of the putative alt-ORF was cloned into a mammalian expression vector with a FLAG-HA epitope tag on the C-terminus of the alt-protein, followed by transfection and immunostaining. As shown in Figure 1e, over-expressed alt-DRAP1 (previously validated^12^ and included as a positive control), alt-PRR3, alt-PRH1, alt-CNPY2 and alt-CACTIN were nucleocytoplasmic. Over-expressed MINAS-60 co-localized with a nucleolar protein, fibrillarin (Figure 1f). Western blotting further confirmed the translation of the six alt-proteins (Figure 1g). MINAS-60 and alt-CNPY2 produced two immunoreactive bands, which is due to multiple in-frame start codon for MINAS-60 (vide infra), and may be caused by multiple start sites, phosphorylation or other post-translational modification for alt-CNPY2, analogous to alt-RPL36^8^.

We hypothesized that the alt-proteins detected specifically under stress conditions are induced in response to cellular stress. To determine whether the expression of alt-CNPY2 is induced by DNA damage, we treated HEK 293T over-expressing alt-CNPY2 with etoposide, followed by western blotting. As shown in Figure 1h, alt-CNPY2 is upregulated up to two fold upon etoposide treatment in both dose- and time-dependent manner. However, the mRNA level of *CNPY2* did not change (Supplementary Figure 1g), indicating that upregulation of alt-CNPY2 is likely due to increased translation or decreased proteolysis, but not increased transcription. Taken together, these results suggest that our chemoproteomic pipeline is able to detect nascent or stress-induced alt-proteins.

### MINAS-60 is cell-cycle regulated and conserved in mouse

We selected MINAS-60 from the BONCAT-detected microprotein dataset for further study because it localizes to the nucleolus, and only one nucleolar microprotein has previously been identified^33^; furthermore, MINAS-60 is nested within the RBM10 CDS, and therefore probing its cellular and molecular roles could shed light on the poorly characterized class of overlapping alt-proteins (Figure 2a, Supplementary Data 2). A different MINAS-60 tryptic peptide was previously detected in human colorectal cancer samples, supporting expression of MINAS-60 in human tissue, but the alt-protein was not defined or characterized in that study^34^. To validate expression of endogenous MINAS-60 from the *RBM10* genomic locus, we generated two independent Cas9-directed knock-in (KI) HEK 293T cell lines with a 3×GFP11-FLAG-HA tag appended to the 3’ end of MINAS-60 alt-ORF^35^, followed with immunostaining. As shown in Figure 2b, endogenously expressed MINAS-60 co-localizes with fibrillarin, consistent with the over-expression results, suggesting MINAS-60 likely functions in the nucleolus.

**Figure 2.**
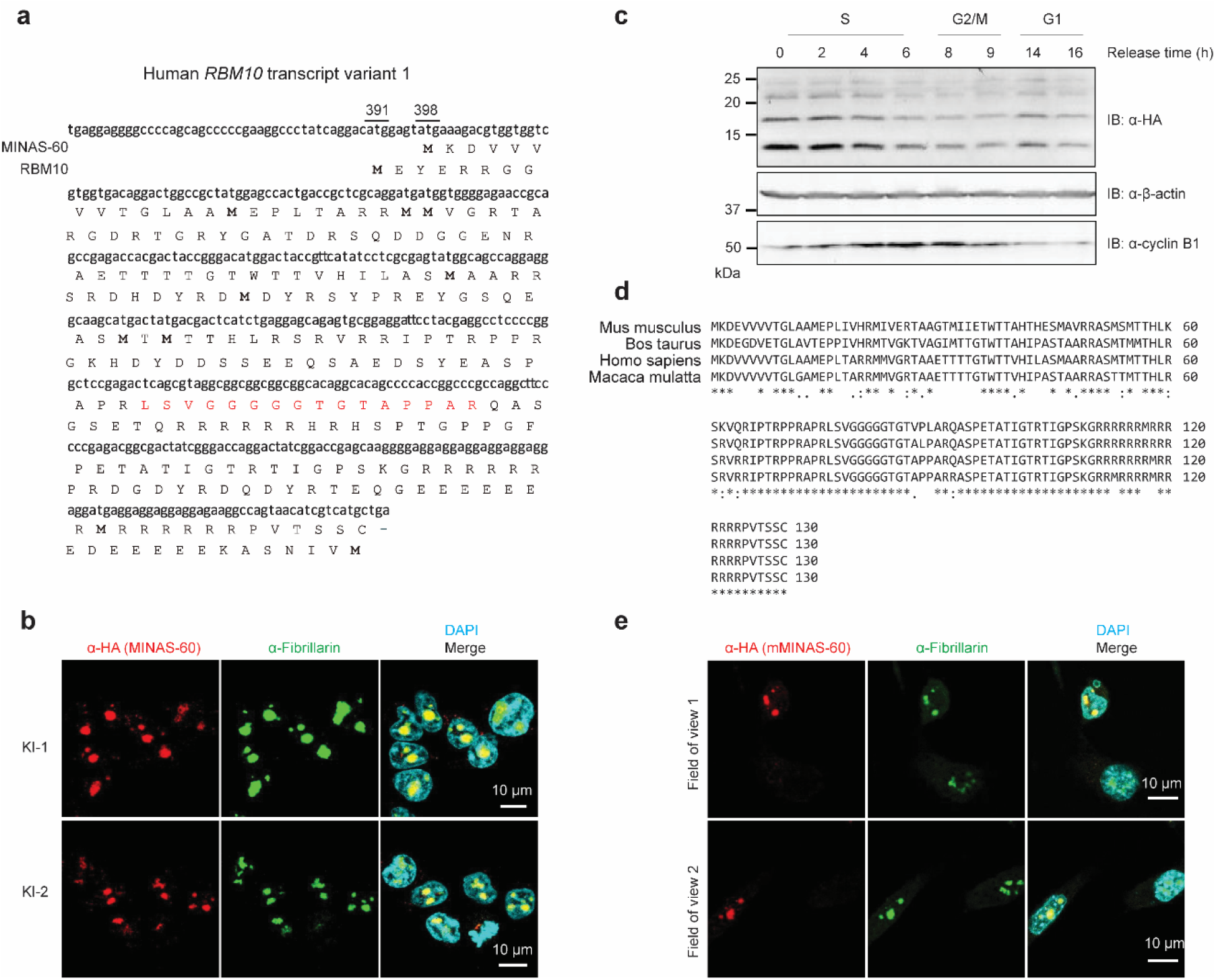
MINAS-60 is endogenously expressed, cell-cycle regulated and conserved in mouse. **a** Shown is the cDNA sequence of human *RBM10* transcript variant 1 with the protein sequences of MINAS-60 and RBM10 indicated below. The start codons of RBM10 (391) and MINAS-60 (398) are numbered above the cDNA sequence. Highlighted in red is the tryptic peptide of MINAS-60 detected by LC-MS/MS in this study. **b** Immunostaining of two MINAS-60 knock-in (KI) HEK 293T cell lines with anti-HA (red), anti-fibrillarin (green), and DAPI (cyan). Scale bar, 10 µm. Data are representative of three biological replicates. **c** Western blotting analysis of synchronized MINAS-60 KI cells released from the G1/S boundary at the indicated time points with antibodies indicated on the right. Data are representative of three biological replicates. **d** RBM10 mRNAs from *Mus musculus* (NM_145627.3), *Bos taurus* (NM_001206693.1), and *Macaca mulatta* (XM_015127275.2) were obtained from NCBI nucleotide database, then translated in the +1, +2 and +3 frames using the Expasy translate tool. Cognate start codons in-frame with sequences homologous to human MINAS-60 were identified in the 5’UTR of each transcript in order to predict the full-length sequence of hypothetical MINAS-60 homologs. Shown is the ClustalW2 alignment of predicted MINAS-60 protein sequences. **e** 3T3 cells transfected with the mouse MINAS-60 (mMINAS-60) expression construct were immunostained with anti-HA (red), anti-fibrillarin (green), and DAPI (cyan). Scale bar, 10 µm. Data are representative of three biological replicates.

The nucleolus is the site of ribosome biogenesis in eukaryotic cells. Ribosome biogenesis starts with the transcription of rDNA, which oscillates during the cell cycle, nearly ceasing during M phase, increasing during G1 phase, and maximizing during S and G2 phases in human cells^36-39^. If MINAS-60 regulates ribosome biogenesis, we hypothesized that the expression of nucleolar MINAS-60 would be correlated with ribosome biogenesis during the cell cycle.

Immunostaining of synchronized MINAS-60 KI HEK 293T cells revealed that nucleolar MINAS-60 staining intensity increased at early S phase, peaking at the 2 h time point. MINAS-60 expression then decreased by late S phase, and was very low during G2/M phase. At G1 phase, the MINAS-60 staining intensity again increased (Supplementary Figure 2). These results were confirmed with western blotting (Figure 2c). MINAS-60 expression is therefore coordinated with ribosome biogenesis activity during the cell cycle.

We then identified the start codon(s) that initiate MINAS-60 translation. Alt-ORFs have been reported to initiate at upstream, non-AUG start codons^8,12,40^ and internal, AUG start codons^7,41^. We tested two upstream in-frame non-AUG start codons, A_383_TC and A_386_GG (numbered relative to the first nucleotide of the cDNA), as well as seven internal AUG start codons (numbered ATG1 – ATG7) (Supplementary Figure 3a). The ∼20 kDa MINAS-60 isoform likely initiates at ATG1 and the ∼15 kDa MINAS-60 isoform likely initiates at ATG6 or 7, because the indicated species is abrogated only when the corresponding start codon is deleted or mutated (Supplementary Figures b-c). These results were further confirmed by over-expressing truncated MINAS-60 coding sequences starting from ATG1 or ATG6 and comparing their products’ sizes with the wild-type construct (Supplementary Figure 3d). The MINAS-60 smORF is therefore entirely contained within the RBM10 coding sequence, and the MINAS-60 ATG1 start codon is only 7 nucleotides downstream of the RBM10 start codon (Figure 2a).

We therefore speculate that MINAS-60 translation may initiate within *RBM10* transcript variant 1 via leaky scanning, and that the additional MINAS-60 isoform diversity observed in KI cells could be generated by alternative splicing, analogous to PTBP3^41^.

ClustalW alignment of hypothetical MINAS-60 homologs from mouse, cattle, and monkey revealed significant sequence similarity, suggesting MINAS-60 is conserved (Figure 2d). To determine whether mouse MINAS-60 is also translated, a mammalian expression vector contains the 5’UTR of mouse *RBM10* transcript variant 1 through the stop codon of the putative mouse MINAS-60 homolog was transfected into 3T3 cells, followed by immunostaining. As shown in Figure 2e, over-expressed mouse MINAS-60 localized to the nucleolus, comparable to human MINAS-60. Taken together, these results indicate that MINAS-60 is endogenously expressed, cell-cycle regulated and conserved from humans to mouse.

### MINAS-60 is associated with nucleolar LSU assembly factors

Because many SEPs characterized to date bind to and regulate other proteins^42^, we performed a co-immunoprecipitation (co-IP) with the nuclear lysates from MINAS-60-FLAG KI cells, and HEK 293T cells as a control. Two major bands were specifically present in the KI co-IP after SDS-PAGE that, upon analysis via label-free quantitative proteomics^43^, yielded 17 proteins enriched >30-fold over control. GO analysis of these 17 hits shows that the top 2 enriched biological processes are ribosome biogenesis and ribosomal large subunit (LSU) biogenesis (Supplementary Figures 4a-c, and Supplementary Data 3). To obtain a more comprehensive picture of MINAS-60-associated proteins, we performed co-IPs and analyzed the entire molecular weight range with quantitative proteomics; while abundant ribosomal proteins limited depth of detection of other proteins, we observed enrichment of four LSU assembly factors: GTPBP4, MRTO4, BRIX1 and NOP2 (Figure 3a and Supplementary Data 4). Co-IP followed by western blotting confirmed the association of these four factors with MINAS-60 (Figure 3b). The association of MINAS-60 with GTPBP4 and MRTO4 did not depend on RNA, because these associations largely survived RnaseA treatment. In contrast, co-purification of MINAS-60 with BRIX1 and NOP2 was severely diminished after treatment with RNaseA, suggesting that their association with MINAS-60 was likely indirect and RNA-dependent (Figure 3b). We therefore hypothesized that MINAS-60 associates with nucleolar, late pre-60S particles containing GTPBP4 and MRTO4 to regulate ribosome biogenesis^18,44^.

**Figure 3.**
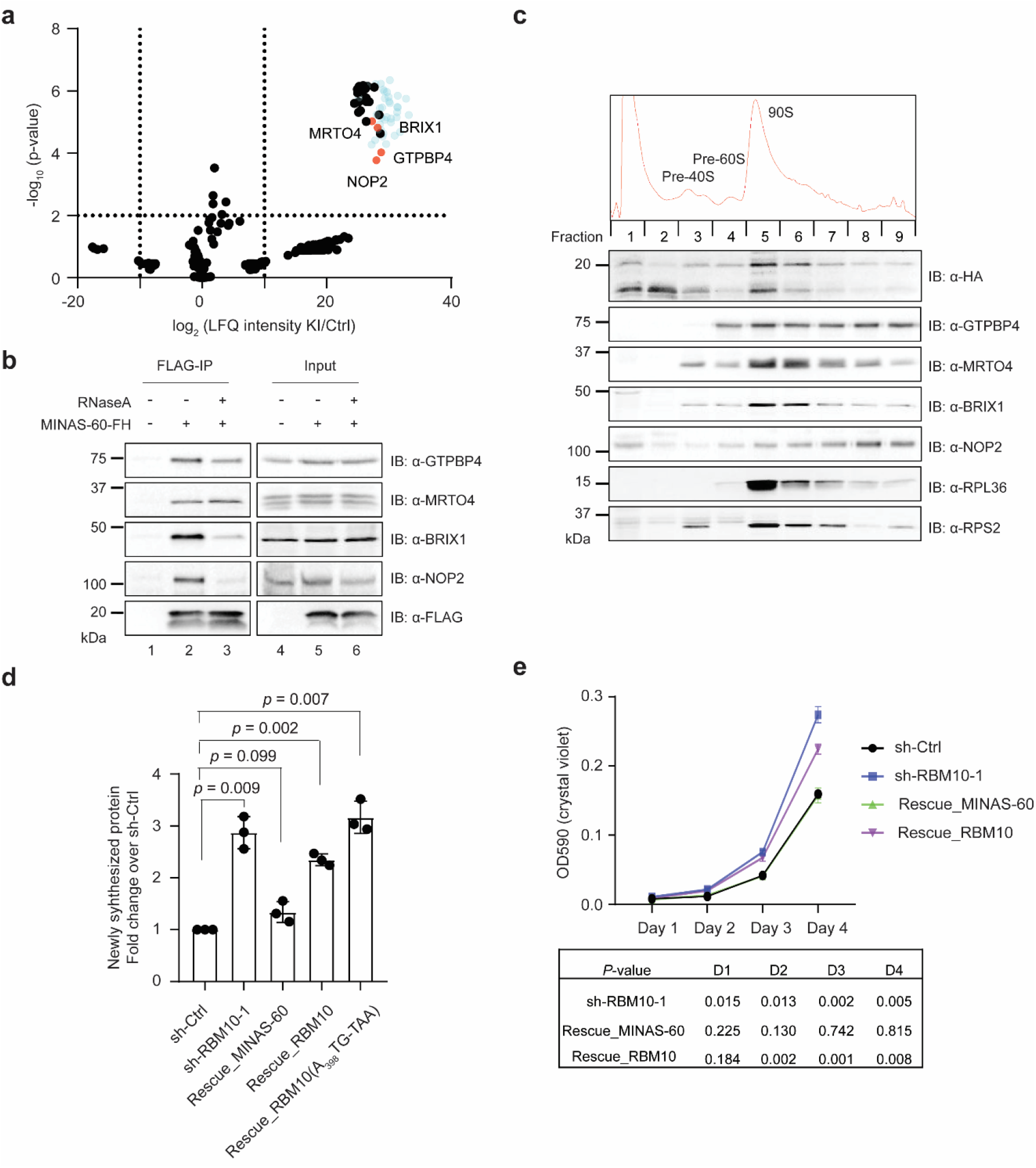
MINAS-60 associates with nucleolar LSU assembly factors, downregulates global protein synthesis and cell proliferation. **a** Volcano plot of quantitative proteomics (*N* = 3) of anti-FLAG pulldown from MINAS-60 KI (KI) or control (Ctrl) HEK 293T nuclear lysates. Ribosomal proteins are indicated in blue. Enriched LSU assembly factors are indicated in red and gene names are labeled. For complete quantitative proteomics results, see Supplementary Data 4. **b** HEK 293T cells were transfected with MINAS-60-FLAG-HA (MINAS-60-FH, lanes 2 and 3) or vehicle (lane 1), and immunoprecipitation (FLAG-IP) was performed in absence (lanes 1 and 2) or presence (lane 3) of RNaseA, followed with immunoblotting (IB) with antibodies indicated on the right. Cell lysates (4%) before IP (input, lanes 4-6) were used as loading controls. **c** Top: Sucrose-gradient sedimentation analysis of nuclear lysates containing ribosome precursor complexes (pre-40S, pre-60S and 90S pre-ribosome) from HEK 293T cells stably expressing MINAS-60-FLAG-HA. Bottom: Western blot analysis of fractions numbered at the top with antibodies indicated on the right. **d** ImageJ was used to quantify the relative puromycin incorporation for cells indicated at the bottom relative to sh-Ctrl from three biological replicates. Data represent mean values ± s.e.m., and significance was evaluated with two-tailed *t*-test. **e** Growth curve of control (sh-Ctrl), *RBM10* knockdown (sh-RBM10-1), rescue with MINAS-60 (Rescue_MINAS-60) and rescue with RBM10 (Rescue_RBM10) HEK 293T cells at the indicated number of days (*N* = 3). Data represent mean values ± s.e.m., and significance was evaluated with two-tailed *t*-test and shown below.

To determine whether MINAS-60 associates with high molecular weight pre-60S particles containing GTPBP4 and MRTO4, we performed sucrose gradient fractionation of nuclear extracts of HEK 293T cells stably expressing epitope-tagged MINAS-60, followed by western blotting analysis. As shown in Figure 3c, a subpopulation of over-expressed MINAS-60 co-sedimented with ribosome assembly factors in high-molecular weight fractions coincident with pre-ribosomal particles. Combined with our co-IP and immunofluorescence data, these results are consistent with a role for MINAS-60 in pre-ribosomal particles.

### MINAS-60 inhibits protein synthesis and cell proliferation

We hypothesized that MINAS-60 could regulate ribosome biogenesis via its association with GTPBP4 and MRTO4. Ribosome biogenesis is required for protein synthesis, which promotes cell growth and proliferation. As a result, ribosome biogenesis is commonly upregulated in cancer cells^21,45^. We reasoned that, if MINAS-60 regulates ribosome biogenesis, its absence should result in changes to protein synthesis and cell proliferation. In order to test this hypothesis, we required a system to query the function of MINAS-60 independent of RBM10, despite their co-encoding on the same transcript. To this end, we knocked down (KD) *RBM10* in HEK 293T cells with two different shRNAs, which silence the entire mRNA and, therefore, both proteins (RBM10 and MINAS-60).

To deconvolute phenotypic effects specific to MINAS-60 in the KD, as well as to exclude off-target effects of shRNA, we generated rescue cell lines stably expressing MINAS-60 (Rescue_MINAS-60) or RBM10 (Rescue_RBM10) on the KD background. qRT-PCR and western blotting analysis revealed that *RBM10* mRNA is efficiently silenced by both shRNAs, and the KD cells successfully re-express MINAS-60 or RBM10 after rescue (Supplementary Figure 5). Noting that the RBM10 rescue construct could be subject to leaky translation to produce both RBM10 and MINAS-60, we also rescued the KD with an RBM10 construct bearing an A_398_TG to TAA mutation that eliminates the first MINAS-60 start codon while preserving RBM10 translation (Supplementary Figure 5).

To test the effect of MINAS-60 expression on cellular protein synthesis, we labeled nascent peptides in the control, RBM10 KD, Rescue_MINAS-60, Rescue_RBM10 and Rescue_RBM10(A_398_TG-TAA) cell lines with puromycin followed by anti-puromycin western blotting^46^. As shown in Figure 3d and Supplementary Figure 6a, depletion of the entire *RBM10* mRNA led to a significant increase in global protein synthesis, and this increase was rescued by reintroduction of MINAS-60. Partial rescue by RBM10 reintroduction was also observed, but was not present in the RBM10(A_398_TG-TAA) rescue cells. Similar results were observed for a second shRNA targeting RBM10 (Supplementary Figures 6b-c). Taken together, these results indicate that MINAS-60 plays an unusual role in ribosome biogenesis to downregulate global protein synthesis. At the same time, RBM10 is unlikely to play a role in this process. We speculate that the partial rescue observed with RBM10 could have been due to leaky expression of MINAS-60 from the internal alt-ORF in the RBM10 construct.

Because protein synthesis, cell growth and proliferation are linked^16,21^, we asked whether MINAS-60 regulates cell proliferation. As shown in Figure 3e, *RBM10* depletion led to a significant increase in cell proliferation, consistent with a published report using HeLa cells^29^. Remarkably, this increase was rescued by reintroduction of MINAS-60 alone, and partially rescued by RBM10 reintroduction. Similar results were observed for a second shRNA targeting RBM10 (Supplementary Figure 6d). These results are consistent with a model in which MINAS-60 inhibits ribosome biogenesis, subsequently downregulating cellular protein synthesis and cell proliferation.

### MINAS-60 inhibits the export of pre-60S ribosome subunits

Eukaryotic ribosome biogenesis can be divided into sequential processes^47^: pre-rRNA transcription; chemical modification and processing of the pre-rRNA, both of which occur in the nucleolus; folding, assembly and maturation of the pre-ribosomal subunits in the nucleolus and nucleus; and export and final maturation of ribosome subunits in the cytoplasm. Because MINAS-60 co-purified with proteins involved in various steps of LSU biogenesis, we wished to determine the step in this process that it regulates.

To determine whether MINAS-60 controls pre-rRNA transcription, we performed qRT-PCR targeting the primary pre-rRNA (47S/45S/30S) using previously published primers^46^ in lysates from control, RBM10 KD, Rescue_MINAS-60 and Rescue_RBM10 cell lines. No significant differences in primary pre-rRNA were observed, suggesting that MINAS-60 does not regulate pre-rRNA transcription (Supplementary Figures 7a-b). To determine whether MINAS-60 regulates the processing of LSU pre-rRNA, we performed northern blotting analysis with the control, RBM10 KD, Rescue_MINAS-60 and Rescue_RBM10 cell lines using a probe complementary to ITS2, which detects all LSU pre-rRNA processing products, including 41S, 32S and 12S pre-rRNAs^46^. As shown in Supplementary Figure 7c, no significant differences were observed between these cell lines, suggesting that MINAS-60 does not regulate the processing of LSU pre-rRNA.

We therefore examined the role of MINAS-60 in LSU assembly and export. To determine whether MINAS-60 regulates nucleocytoplasmic export of the pre-60S ribosomal subunit, we quantified the ratio of nuclear vs. cytoplasmic RPL29-GFP^48^ stably expressed in the control, RBM10 KD, Rescue_MINAS-60 and Rescue_RBM10 cell lines. As shown in Figure 4a and Supplementary Figure 8a, depletion of RBM10 decreased the ratio of nuclear to cytoplasmic RPL29-GFP, which can be rescued by reintroduction of MINAS-60, and partially rescued by RBM10 reintroduction, likely due to leaky MINAS-60 translation. Similar results were observed for a second shRNA targeting RBM10 (Figure 4a, right, and Supplementary Figures 8b-c). These observations are consistent with the inhibition of cytoplasmic export of pre-60S subunits by MINAS-60. As a control, to determine whether MINAS-60 regulates nucleocytoplasmic export of the pre-40S ribosomal subunit, we quantified the ratio of nuclear vs. cytoplasmic RPS2-GFP^48^ stably expressed in the control, RBM10 KD, Rescue_MINAS-60 and Rescue_RBM10 cell lines. As shown in Supplementary Figure 9, no significant changes were observed between these cell lines, suggesting MINAS-60 does not regulate the assembly or export of 40S ribosomal subunits.

**Figure 4.**
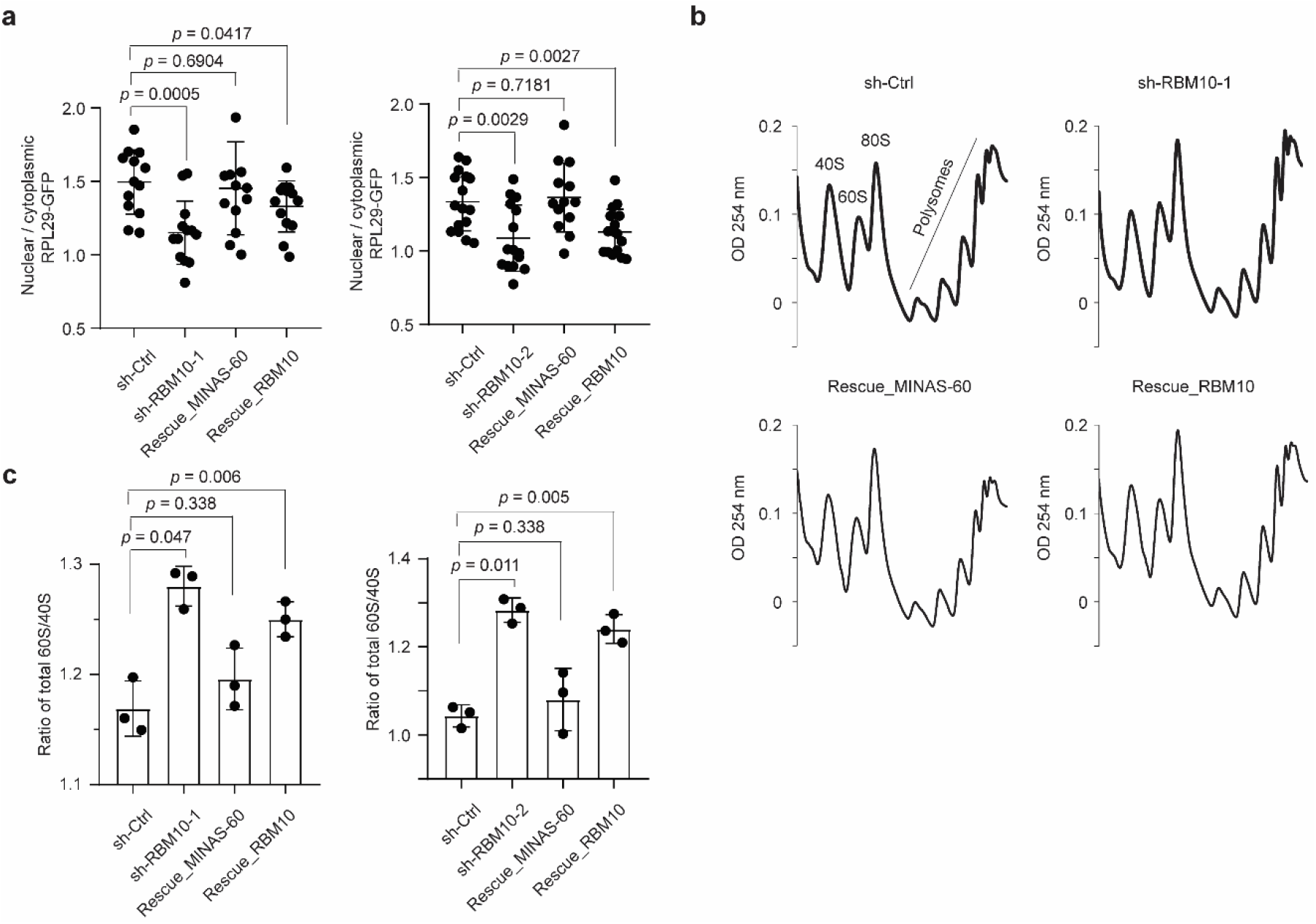
MINAS-60 inhibits LSU export. **a** Quantitation of the ratio of RPL29-GFP intensity in the nucleus vs. cytoplasm in control (sh-Ctrl), *RBM10* knockdown with one of the two shRNAs (sh-RBM10-1 (left), sh-RBM10-2 (right)), rescue with MINAS-60 (Rescue_MINAS-60), or rescue with RBM10 (Rescue_RBM10) HEK 293T cells stably expressing RPL29-GFP. At least 13 fields of view were analyzed, totaling > 350 cells for each measurement. Data represent mean values ± s.e.m., and significance was evaluated with two-tailed *t*-test. **b** Sucrose-gradient sedimentation analysis of ribosomal fractions (40S, 60S, 80S and polysomes) of cytoplasmic lysates from control (sh-Ctrl), *RBM10* knockdown (sh-RBM10-1), rescue with MINAS-60 (Rescue_MINAS-60) or rescue with RBM10 (Rescue_RBM10) HEK 293T cells. Data are representative of three biological replicates. **c** Quantitation of the ratio of cytoplasmic 60S to 40S subunits in the cell lines indicated below after sucrose gradient fractionation. The area under each peak was measured using ImageJ following a previously published method^61^. Data represent mean values ± s.e.m., and significance was evaluated with two-tailed *t*-test.

The observation of upregulated pre-60S export in the absence of MINAS-60 predicted that the same conditions should lead to an increase in mature cytoplasmic 60S ribosomal subunits. To test this, we performed cytoplasmic polysome profiling. As shown in Figures 4b-c, knockdown of RBM10 increased the ratio of cytoplasmic 60S/40S ribosome subunits, and this increase can be rescued by reintroduction of MINAS-60, and partially rescued by RBM10. Similar results were observed for a second shRNA targeting RBM10 (Figure 4c, right and Supplementary Figure 10). Taken together, these results suggest that MINAS-60 specifically decreases mature large ribosomal subunits by negatively regulating the assembly or export of pre-60S particles.

### MINAS-60 inhibits the late-stage assembly of pre-60S

Lastly, we speculated that MINAS-60 functions as a checkpoint inhibitor in pre-60S assembly prior to export from the nucleus, which would suggest that the protein composition of LSU precursors should change in cells lacking MINAS-60. MRTO4 and BRIX1 are LSU biogenesis factors present in multiple intermediate-to-late, or early-to-intermediate, pre-60S particles, respectively, and we examined their interactomes as a readout of changes in pre-60S protein composition in the presence or absence of MINAS-60 (Figure 5a)^18^. We stably expressed MRTO4-FLAG or BRIX1-FLAG, in control or RBM10 KD HEK 293T cells to enable affinity purification of pre-ribosomal particles, followed by quantitative proteomics and western blotting. We detected statistically significant increases in several late LSU assembly factors co-purified with MRTO4 in RBM10 KD cells (Figure 5b, Supplementary Data 5). These increases were further confirmed by western blotting (Figure 5c). Similar results were observed for BRIX1 co-IP, though fold changes were smaller (Figures 5d-f, Supplementary Data 6). These results suggest that remodeling of pre-60S particles toward more mature stages occurs in the absence of MINAS-60, consistent with MINAS-60 acting as an inhibitor for LSU assembly and export.

**Figure 5.**
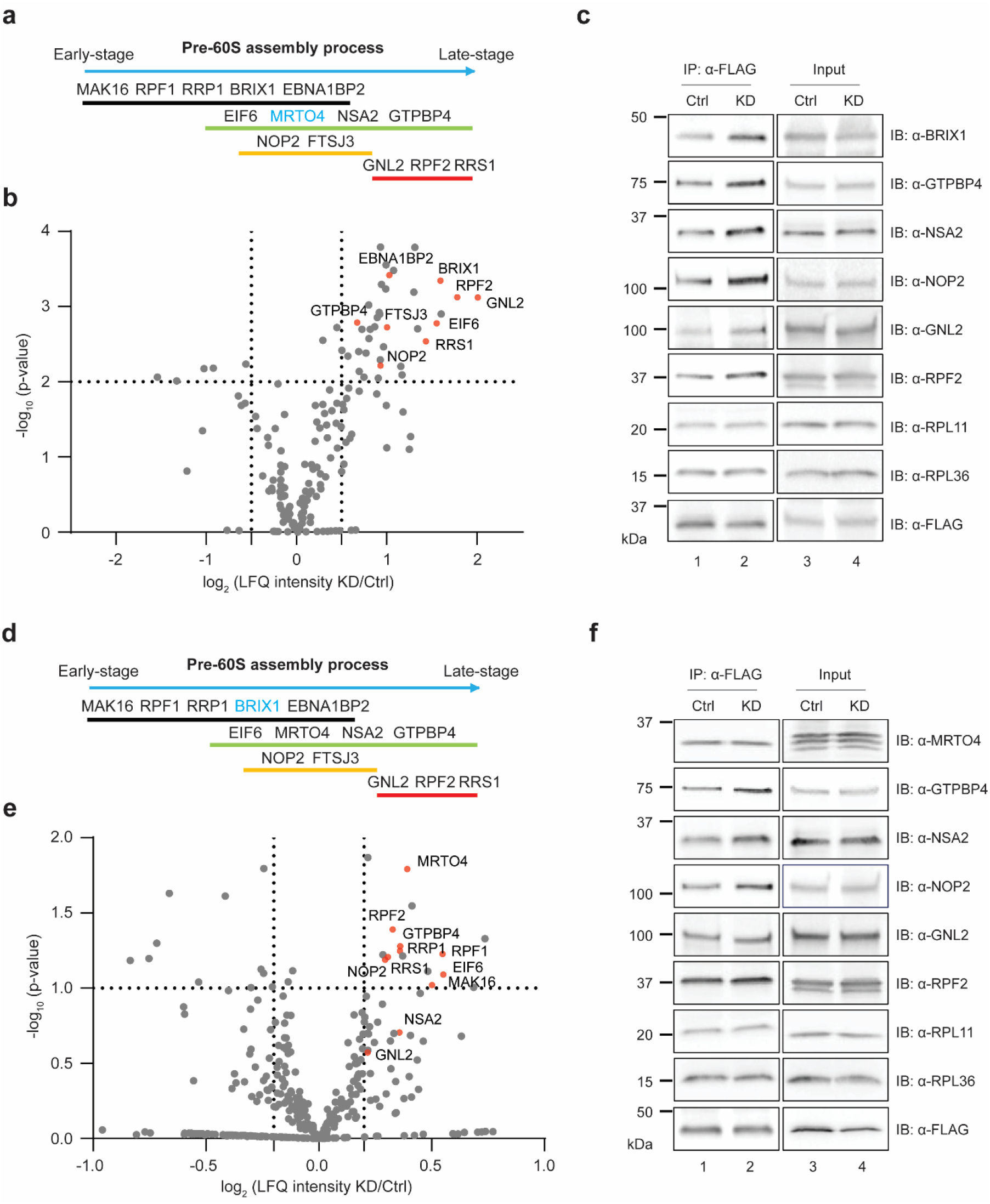
*RBM10* silencing promotes pre-60S assembly. **a, d** Schematic representation of pre-60S assembly factors associated with different states based on a pre-60S structure report^18^. The bait protein MRTO4 (**a**) and BRIX1 (**d**) is indicated in blue. **b, e** Volcano plot of quantitative proteomics (*N* = 4 (**b**), *N* = 5 (**e**)) of MRTO4-FLAG (**b**) or BRIX1-FLAG (**e**) pulldown from HEK 293T cells stably expressing the bait protein and control shRNA (Ctrl), or the bait protein and *RBM10* shRNA (KD), to quantify relative changes in the bait protein co-IP of LSU assembly factors in *RBM10* KD over control HEK 293T cells. Increased assembly factors are indicated in red and gene names are labeled. For complete quantitative proteomics results, see Supplementary Datas 5 and 6. **c, f** MRTO4 FLAG-IP (**c**) or BRIX1-FLAG-IP (**f**) and western blotting with antibodies indicated on the right using the two cell lines described above. Cell lysates (4%) before IP (input) were used as the loading control. Data are representative of three biological replicates.

## Discussion

In this work, we developed a bio-orthogonal strategy for the direct detection of unannotated nascent or stress-induced alt-proteins. BONCAT has been powerfully applied in prior studies to examine nascent protein synthesis, particularly in neurons^30^, but standard BONCAT workflows, like many proteomics protocols, include column or gel resolution steps that de-enrich small proteins and peptides, and are therefore refractory to detection of microproteins and alt-proteins. For the first time in this work, we followed azidohomoalanine labeling with small protein enrichment using a previously reported C8 column strategy^31^, coupled with direct on-bead capture to eliminate the need for column-based removal of excess biotin probe molecules, which also remove small proteins.

Interfacing this BONCAT-based chemoproteomic pipeline with our platform technology for unannotated microprotein and alt-protein detection, we identified 22 alt-proteins undergoing active synthesis in HEK 293T cells. For six selected alt-proteins, we verified their translation, one of which, alt-CNPY2, is likely post-transcriptionally upregulated by DNA damage stress. We furthermore propose that BONCAT-mediated detection of MINAS-60 reflects the subpopulation of cells that actively synthesize this protein during S phase of the cell cycle, consistent with its coordinated synthesis with the ribosome biogenesis machinery. Taken together, these results show that our method is able to detect unannotated alt-proteins expressed during the cell cycle and cellular stress, and that these proteins may play important roles in the cell. In the future, an important next step will be to characterize the biological role of alt-CNPY2, and to expand this strategy to additional cellular stress conditions.

We then conducted a focused functional study of nucleolar MINAS-60, loss of which promotes the late-stage assembly of pre-60S and the export of pre-60S particles into cytoplasm, leading to increases in cytoplasmic 60S ribosomal subunits, global protein synthesis, and cell proliferation (Figure 6). The MINAS-60 alt-protein is therefore a repressor of LSU biogenesis. In contrast to smORF-encoded proteins, few alt-ORFs nested within canonical protein CDS have been defined in molecular detail. This work, combined with previous literature^7,49^, expands the recent finding that a single human transcript can encode overlapping, sequence-independent, yet functionally related proteins.

**Figure 6.**
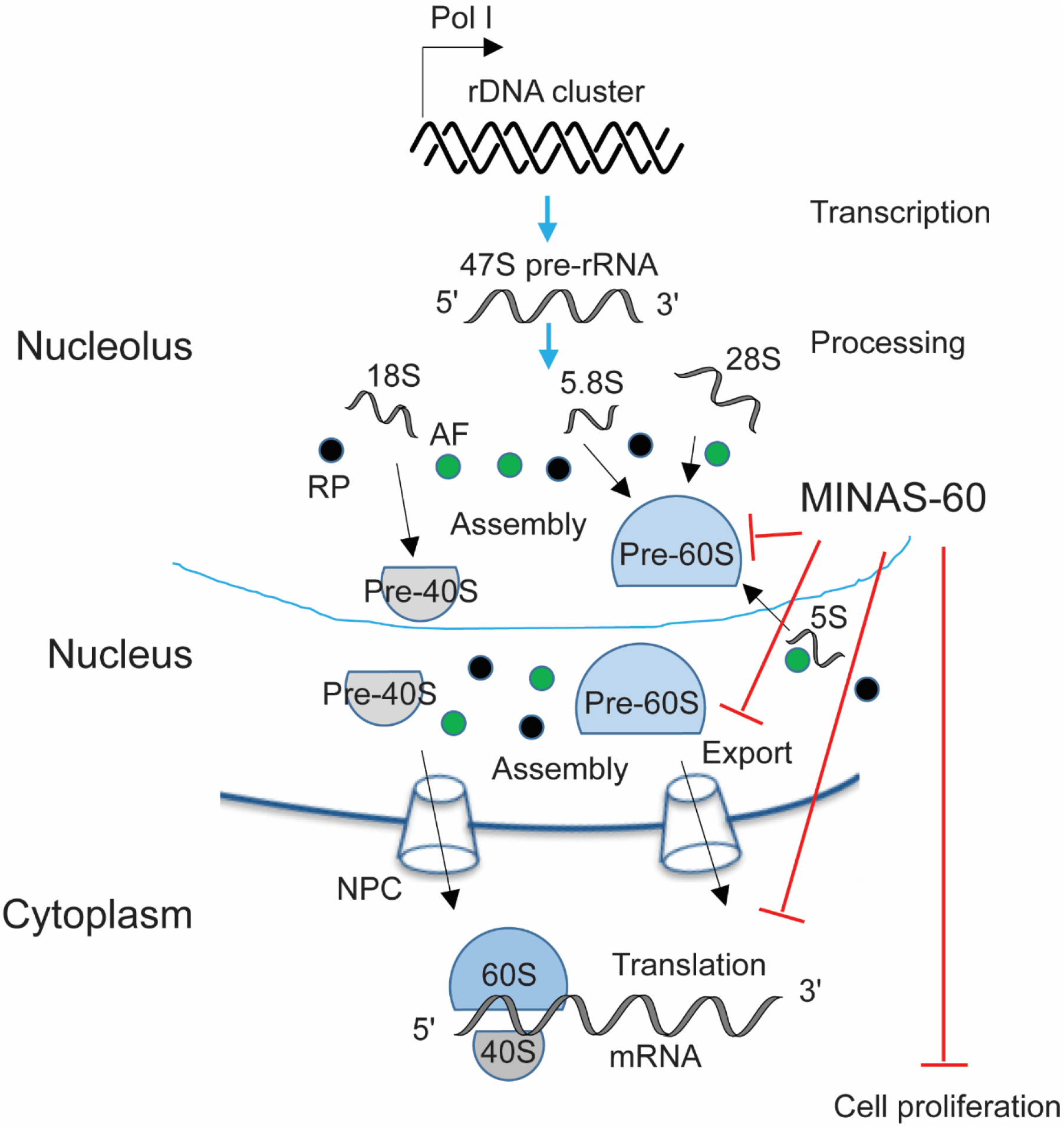
Model of MINAS-60 regulatory pathway. MINAS-60 localizes to the nucleolus, where it associates with multiple pre-60S assembly factors, including GTPBP4 and MRTO4, to inhibit the late-stage pre-60S assembly and the export of pre-60S into cytoplasm, consequently downregulating global protein synthesis and cell proliferation.

Importantly, the *RBM10* gene plays important roles in human physiology and disease. RBM10 (RNA Binding Motif 10) is an RNA binding protein that regulates alternative pre-mRNA splicing^29,50^. Null mutations in the *RBM10* gene are found in patients with TARP syndrome^51^, an X-linked inherited pathology associated with malformation of multiple organs and significant early-life mortality. The *RBM10* gene was also found to be among the most frequently mutated genes in lung adenocarcinoma samples, and RBM10 inhibits cancer cell proliferation^52^.

The finding that *RBM10* dually encodes MINAS-60, which also downregulates cell proliferation via repression of ribosome biogenesis, opens the question of whether this alt-protein contributes to *RBM10* mutation associated disease phenotypes, analogous to alt-FUS, which forms pathogenic cytoplasmic aggregates similar to those caused by the co-encoded FUS protein^3^.

Several studies have suggested that as many as 50% of multicistronic human genes encode proteins that directly interact to form complexes^9,53^. In contrast, the alt-protein alt-RPL36 can regulate the same pathway (translation) as the canonical protein co-encoded with it (RPL36), without directly interacting with each other^8^. In this work, we show that MINAS-60 and RBM10 exhibit comparable antiproliferative cellular effects but act via independent pathways: ribosome biogenesis vs. alternative pre-mRNA splicing, respectively. While it remains possible that RBM10 regulates splicing of some nucleolar proteins, RBM10 reintroduction did not rescue the protein translation defect observed in cells lacking *RBM10* expression in this study. Based on these results we conclude that RBM10 does not significantly contribute to ribosome biogenesis. We therefore speculate that selective pressure for co-encoding multiple functional proteins in multicistronic human genes may act at the level of cellular fitness, and direct interaction between alt-proteins and canonical proteins is only one mechanism among many by which the fitness effects of co-encoded proteins may be optimized. Another possible mechanism is that, since alt-ORFs may represent protogenes^54^, *de novo* acquisition of a novel alt-ORF within a protein coding gene might lead to positive selection if both the alt-ORF and canonical protein coding sequence increase cellular fitness under the same conditions, without necessarily acting on the same pathway or process. This broadens the current models of multicistronic human gene function^9^.

To our knowledge, MINAS-60 is only the second human microprotein or alt-protein validated to localize to the nucleolus to date, and the first reported to regulate ribosome biogenesis. A previous study identified nucleolar microprotein C11orf98, which interacts with nucleolar proteins NPM1 and NCL, but no role for this factor in ribosome biogenesis has been investigated^33^. In this work, we reported MINAS-60 as the first protein that specifically inhibits LSU biogenesis, and we suspect that it functions as a mammal-specific checkpoint to ensure correct pre-60S assembly prior to nuclear export. Supporting the hypothesis that MINAS-60 has arisen in higher eukaryotes, the *RBM10* gene is not conserved in yeast, though we cannot yet rigorously conclude that a functional homolog of MINAS-60 is absent in lower eukaryotes. Direct experimental evidence will be required to test the hypothesis that MINAS-60 could represent a control point for ribosome biogenesis that is unique to mammalian cells.

Interestingly, another mammal-specific, putative LSU biogenesis inhibitor has recently been identified. A structural study revealed that an unidentified protein (protein X) is positioned to block the incorporation of nuclear export factor NMD3 into the late pre-60S particle, hypothetically suppressing pre-60S export into the cytoplasm^44^. Protein X binds in the immature peptidyl transfer center and directly contacts GTPBP4 as well as helix 89 of the rRNA. Importantly, protein X was specifically observed in cryo-electron microscopy structures of human pre-60S particles and has not been detected in structures of yeast LSU precursors^44^.

Based on the observations that 1) both MINAS-60 and protein X likely function as repressors of LSU biogenesis at the late assembly and export stage; and 2) both MINAS-60 and protein X associate with GTPBP4, we speculate that MINAS-60 functions by a similar mechanism to protein X. It is possible that protein X and MINAS-60 are the same protein, and that protein X could not be identified due to its absence from the UniProt database; however, efforts to model MINAS-60 sequences into the reported structures remain inconclusive to date. It is therefore also possible that multiple previously unknown mammalian proteins regulate LSU biogenesis, suggesting that regulation of ribosome biogenesis may be more complex in human cells than in yeast.

Why, then, might MINAS-60 have arisen in mammalian cells? In order to achieve a balance between cellular growth requirements and energy-intensive ribosome production^15^, and to ensure that improperly assembled ribosome subunits do not lead to mistranslation^23^, ribosome biogenesis needs to be precisely monitored in cells. For example, the Rio1-Nob1-Pno1 network establishes a checkpoint to safeguard against the release of immature 40S subunits into translating ribosomes^55^. However, while hundreds of proteins required for ribosome biogenesis in human cells have now been identified^56^, few repressors of ribosome biogenesis have been reported, and the ones that have been identified predominately act on pre-ribosomal RNA transcription^57-59^. LSU biogenesis involves multiple composition and conformational changes during generation of the peptidyl transferase center (PTC). Our data lead us to speculate that MINAS-60 and/or Protein X may act as checkpoints for this process to safeguard cellular energy expenditure and/or faithful PTC formation. This hypothesis could be tested by examining the effects of MINAS-60 on cellular fitness under nutrient stress, as well as on the fidelity and accuracy of global protein translation.

In conclusion, this study, combined with previous literature^13^, demonstrates the power of chemoproteomics to reveal alt-proteins that are regulated and/or functional in important cellular processes including DNA damage and ribosome biogenesis, and reveals an entirely new regulatory node of the ribosome biogenesis pathway, meriting further development of chemical tools to enrich and identify functional alt-proteins.

## Online methods

### Data analysis

Two-tailed *t*-test were performed using Microsoft Excel or GraphPad Prism, and *F*-tests were performed to evaluate equal variance between samples.

### Antibodies

Primary antibodies for western blotting include the following: anti-FLAG (Sigma, F3165 or Cell Signaling, 14793); anti-HA (Invitrogen, 71-5500); anti-β-actin (Invitrogen, BA3R); anti-GTPBP4 (Abclonal, A4565); anti-MRTO4 (ThermoFisher, 20194-1-AP); anti-NSA2 (Abclonal, A14475); anti-NOP2 (Cell Signaling, 25017); anti-BRIX1 (Abclonal, A14481); anti-RPF2 (Abclonal, A17224); anti-GNL2 (Abclonal, A13191); anti-RPL11 (Cell Signaling, 18163); anti-RPL36 (Bethyl Laboratories, A305065A-M); anti-RPS2 (Invitrogen, PA5-30160); anti-RBM10 (Abcam, ab72423); anti-puromycin (Kerafast, EQ0001); anti-GFP (Abcam, ab183734); anti-cyclin B1 (Cell Signaling, 4138).

Immunoprecipitation was performed with anti-FLAG M2 affinity gel (Sigma, A2220). Secondary antibodies for western blotting are goat anti-rabbit IgG horseradish peroxidase conjugate (Rockland, 611-1302) and goat anti-mouse IgG horseradish peroxidase conjugate (Rockland, 610-1319-0500). Primary antibodies for immunostaining are rabbit anti-HA (Invitrogen, 71-5500) and mouse anti-Fibrillarin (abcam, ab4566). Secondary antibodies for immunostaining are goat anti-rabbit IgG Alexa fluor 568 (Invitrogen, A11011) and goat anti-mouse IgG Alexa fluor 647 (Invitrogen, A21235).

### Cloning and genetic constructs

A DNA sequence comprising the full 5’UTR of human *DRAP1* transcript, *PRR3* transcript variant 2, *PRH1* transcript variant 1, *CNPY2* transcript variant 1, *CACTIN* transcript variant 1, or *RBM10* transcript variant 1 through the stop codon of the relative alt-protein was amplified by PCR with a dual FLAG and HA epitope tag appended to the 3’ end of the alt-protein coding sequence from an in-house library of reverse-transcribed HEK 293T cDNAs, then subcloned into pcDNA3. Deletion or mutation constructs of MINAS-60 bearing a dual FLAG and HA tag were generated by ligating PCR products into BamHI and EcoRI cloning sites in the pcDNA3 vector. For generation of HEK 293T cells stably expressing MINAS-60, a dual FLAG and HA tag were appended to the 3’ end of MINAS-60 by PCR, and the dually tagged coding sequence was then cloned into pLJM1. The cDNA clone expressing RPS2 was purchased from Addgene (a gift from [Thomas Tuschl](https://www.addgene.org/Thomas_Tuschl/)), and the coding sequences of MRTO4, BRIX1 and RPL29 were amplified by PCR from an in-house HEK 293T cDNA library, then subcloned into pJLM1 for producing lentivirus. RNAi constructs were made by synthesizing oligonucleotides encoding a 21 bp short hairpin RNA that targets RBM10 (shRNA1, CTTCGCCTTCGTCGAGTTTAG; shRNA2, TCCAACGTGCGCGTCATAAAG), then subcloned into pLKO.1. The empty pLKO.1 vector control was purchased from Sigma (SHC001). qPCR primer sequences are provided in Supplementary Table 2.

### Cell culture, lentivirus production and stable cell line generation

HEK 293T cells were purchased from ATCC and early-passage stocks were established in order to ensure cell line identity. Cells were maintained up to only 10 passages. HEK 293T cells were cultured as previously described^8^. To produce lentivirus and generate stable cell lines, HEK 293T cells were co-transfected using polyethyleneimine (Polysciences, 23966) with expression construct in pLJM1, along with pMD2.G and psPAX2, and growth media were replaced after 7-8 h. 48 h post-transfection, media containing viruses was harvested, filtered through a 0.45-μm filter, and infection was performed by mixing with two volumes of fresh media containing suspended HEK 293T cells. 24 h post-infection, the growth media was replaced. 48 h post-infection, stably expressing cells were selected with 4 µg/mL puromycin for 2 days. Early stocks of stable cell lines were established after selection. Stable cell lines were released from puromycin for 2 days prior to use in experiments.

### Immunostaining and live-cell imaging

HEK 293T cells were plated on glass coverslips and transfected the next day. Forty-eight hours later, the cells were fixed in 10% formalin for 15 min at room temperature (RT), permeabilized with PBS containing 0.2% (v/v) TritonX-100, then incubated with primary antibodies for 18 h at 4°C. After washing with PBS, the cells were incubated with secondary antibodies and DAPI for 1 h at RT, washed with PBS and mounted with Mowiol (Sigma, 81381) before viewing.

HEK 293T cells stably expressing RPL29-GFP or RPS2-GFP were grown to 70% confluency on coverslips in 12-well plates. Coverslips were inverted and imaged in pre-warmed DMEM with 10% FBS, 1% penicillin-streptomycin in MatTek imaging dishes. Confocal imaging was performed on a Leica SP8 LS confocal microscope with 63× oil immersion objective with atmosphere-controlled stage at 37°C. Nuclear/cytoplasmic ratios of RPL29 and RPS2 were measured using the ImageJ Intensity Ratio Nuclei Cytoplasm Tool (RRID:SCR_018573; https://github.com/MontpellierRessourcesImagerie/imagej_macros_and_scripts/wiki/Intensity-Ratio-Nuclei-Cytoplasm-Tool).

### Immunoprecipitation and proteomics

Control HEK 293T cells or MINAS-60 KI cells were grown to 80-90% confluency in 15 cm dishes. Cells were harvested and suspended in 1 mL nuclear isolation buffer (10 mM HEPES-KOH pH 7.4, 100 mM KCl, 5 mM MgCl_2_ with 0.5% NP40 and Roche Complete protease inhibitor cocktail tablets (Roche, 11873580001)), and incubated on ice for 10 min, followed by centrifugation at 3,000 g, 4°C, 3 min. The nuclear pellets were suspended in 1 mL lysis buffer (Tris-buffered saline (TBS) with 1% Triton X-100 and Roche Complete protease inhibitor cocktail tablets), followed with sonication and immunoprecipitation as previously described^8^. After the final wash, elution was in 40 µL of 3× FLAG peptide (Sigma, F4799), at a final concentration of 100 µg/mL in lysis buffer at 4°C for 1 h. The eluted proteins were subjected to SDS-PAGE separation prior to LC-MS/MS analysis.

Gel slices, containing either resolved protein bands or entire lanes, were digested with trypsin at 37°C for 14-16 h. The resulting peptide mixtures were extracted from the gel, dried, subjected to ethyl acetate extraction to remove residual detergent, de-salted with peptide cleanup C18 spin column (Agilent Technologies, 5188-2750), then re-suspended in 35 µL 0.1% formic acid (FA), followed by centrifugation at 21,130 g, 4°C, 30 min. A 5 μL aliquot of each sample was injected onto a pre-packed column attached to a nanoAcquity UPLC (Waters) in-line with a Thermo Scientific™ Q Exactive™ Plus Hybrid QuadrupoleOrbitrap™ mass spectrometer (Thermo Scientific) and a 130-min gradient was used to further separate the peptide mixtures as follows (solvent A: 0.1% FA; solvent B: acetonitrile (ACN) with 0.1% FA): Isocratic flow was maintained at 0.1 μL/min at 1% B for 40 min, followed by linear gradients from 1% B to 6% B over 2 min, 6% B to 24% B over 48 min, 24% B to 48% B over 5 min, 48% B to 80% B over 5 min. Isocratic flow at 80% B was maintained for 5 min, followed by a gradient from 80% B to 1% B over 5 min, and isocratic flow at 1% B was maintained for 10 min. The full MS was collected over the mass range of 300-1,700 m/z with a resolution of 70,000 and the automatic gain control (AGC) target was set as 3 × 10^6^. MS/MS data was collected using a top 10 high-collisional energy dissociation method in data-dependent mode with a normalized collision energy of 27.0 eV and a 1.6 m/z isolation window. MS/MS resolution was 17,500 and dynamic exclusion was 90 seconds.

For identification of alt-and microproteins, ProteoWizard MS Convert was used for peak picking and files were analyzed using Mascot. Oxidation of methionine and N-terminal acetylation were set as variable modifications, and a previously reported^12^ three-frame translation of assembled transcripts from HEK 293T mRNA-seq was used as the database. For co-IP proteomics searches and quantitative analysis, files were analyzed using MaxQuant, oxidation of methionine and N-terminal acetylation were set as variable modifications, and human UniProt plus MINAS-60 was used as the database for searching. For all analysis, a mass deviation of 20 p.p.m. was set for MS1 peaks, and 0.6 Da was set as maximum allowed MS/MS peaks with a maximum of two missed cleavages. Maximum false discovery rates (FDR) were set to 1% both on peptide and protein levels. Minimum required peptide length was five amino acids.

Protein quantitation was accomplished by calculating the LFQ intensity ratio of KI or KD pulldown to negative control samples using MaxQuant (version 1.6.8.0) with standard parameters.

### Dibenzocyclooctyne (DBCO) bead construction

NHS-activated beads (Pierce, 88826) were conjugated to dibenzocyclooctyne-amine (Sigma, 761540) by 90 min incubation at room temperature with rotation in a saturated 100 mM sodium bicarbonate (pH 8.0) solution. The beads were then washed and blocked for 2 h with 3 M ethanolamine according to the manufacturer’s instructions. After blocking, the beads were re-suspended in PBS prior to immediate protein conjugation.

### BONCAT (bio-orthogonal non-canonical amino acid tagging)

HEK 293T cells were grown to 80-90% confluency in 15 cm dishes, treated with the methionine aminopeptidase inhibitor TNP470 (50 nM) for 2 h, then immersed in methionine-free DMEM (Corning, 17-204-CI) for 30 min before addition of 4 mM AHA (Click Chemistry tools, 1066-1000) in methionine-free DMEM with 10% FBS. For stress conditions, cells were treated with 4 mM AHA with simultaneous exposure to 500 µM sodium arsenite, 20 µM etoposide (Sigma, E1383), or 1 mM DTT. After a 2 h incubation, the cells were washed twice with cold PBS, harvested, and flash frozen until further processing.

### DBCO bead enrichment of AHA-labeled proteins and ERLIC (electrostatic repulsion hydrophilic interaction chromatography) fractionation

AHA-labeled cells were lysed by boiling in 50 mM HCl with 0.01% 2-mercaptoethanol and 0.05% TritonX-100 for 10 min. The cells were then pelleted at 21,100 g, 4°C for 30 min, and passed through a 5 µm filter before size selection with a C8 column (Agilent Technologies, 12102100) essentially as previously reported^32^.

The C8 column was pre-conditioned with 1 bed volume of methanol and 2 bed volumes of 0.25 M triethylammonium formate (TEAF, pH 3.0). Then up to 2 bed volumes of cell lysate were loaded on the column, followed by twice washes with 2 bed volumes of TEAF and elution with 2 bed volumes of ACN:TEAF (1:3). The sample was then dried and reconstituted in 300 µL PBS. Cell lysates were incubated with DBCO beads for 1 h at RT. The beads were washed twice with 1 mL RIPA buffer, once with 1 M KCl, once with 0.1 M sodium carbonate, once with 2 M urea, twice with RIPA buffer, and finally 6 times with PBS. Proteins covalently conjugated to the beads were then subjected to reduction, alkylation and on-bead trypsin digestion according to standard protocols^62^.

Before LC-MS/MS, the digested peptides were fractionated using ERLIC on an Agilent 1100 HPLC. Peptides were re-suspended in 55 µL of 85% ACN/0.1% FA and 50 µL was loaded onto a polyWAX LP column (150×1.0 mm; 5 µm 300 Å; PolyLC). Samples were run on an 80 min gradient protocol as follows (Solvent A: 80% ACN 0.1% FA; Solvent B: 30% ACN 0.1% FA): Isocratic flow was maintained at 100% A at a flow rate of 0.3 mL/min for 5 min, followed by a 17 min linear gradient to 8% B, and a 25 min linear gradient to 45% B. Finally, a 10 min gradient to 100% B was followed by a 5 min hold at 100% B before a 10 min linear gradient back to 100% A, followed by an 8 min hold at 100% A. The digested peptides were separated into 12-15 fractions which were dried and re-suspended in 7 µL of 3:8 70% FA: 0.1% TFA before LC-MS/MS analysis.

### Generation of knock-in cell lines

MINAS-60 3xGFP11-FLAG-HA KI HEK 293T cells were generated using CRISPR-Cas9. Guide RNAs (gRNAs) were designed with the guide design tool from the Zhang lab (crispr.mit.edu) to target the RBM10 genomic region gRNA1, 5’-TGTCGGCCAGGATTCCTACG-3’; gRNA2, 5’-CCCGATAGTCGCCGTCTCGG-3’. Double-stranded DNA oligonucleotides corresponding to the gRNAs were inserted into pSpCas9(BB)-2A-GFP vector (Addgene, as a gift from F. Zhang, MIT, Cambridge, MA). A donor plasmid containing 300 bp homology left-arm and 300 bp homology right-arm sequence around the stop codon of MINAS-60, which are separated with 3xGFP11-FLAG-HA tag and BamHI / NotI restriction sites was synthesized by GenScript, and a DNA sequence containing pGK promoter and hygromycin resistance gene were subcloned into the donor plasmid using the BamH1 and NotI restriction sites. An equal mixture of the gRNA and donor plasmids were transfected into HEK 293T cells using polyethyleneimine, and hygromycin selection was performed 2 days post-transfection. MINAS-60-3xGFP11-FLAG-HA KI cells were confirmed by genomic DNA PCR and sequencing.

### Puromycin incorporation

SUnSET was used to measure protein synthesis^63^. Briefly, HEK 293T cells were grown to 80-90% confluency in 6 well plates, then growth media was replaced with media containing 1 μM puromycin and cells were incubated at 37°C in a humidified atmosphere with 5% CO_2_ for 1 h according to a previously published protocol^56^. The cells were then washed once with PBS, harvested and analyzed with western blotting.

### Sucrose gradient profiling

HEK 293T cells were seeded in 15 cm dishes at 2.5×10^7^ cells per dish and cultured 24 h. Cells were then treated with 100 µg/mL cycloheximide for 5 min, washed with cold PBS containing 100 µg/mL cycloheximide twice, harvested and flash frozen until further processing.

For cytoplasmic polysome profiling, after thawing on ice, cells were lysed in polysome lysis buffer (20 mM HEPES-KOH pH 7.4, 100 mM KCl, 5 mM MgCl_2_, 100 µg/mL cycloheximide, 1 mM DTT with 1% TritonX-100, Roche Complete protease inhibitor cocktail tablets and Ribonuclease Inhibitors (Promega N2511)), and incubated on ice for 10 min, followed by centrifugation at 21,130 g, 4°C, 10 min. The supernatants were normalized according to absorbance (A260) and layered onto 12 mL 10-50% sucrose gradients (20 mM HEPES-KOH pH 7.4, 100 mM KCl, 5 mM MgCl_2_, 100 µg/mL cycloheximide, 1 mM DTT with Roche Complete protease inhibitor cocktail tablets and Ribonuclease Inhibitors), followed with centrifugation in an SW-41Ti rotor at 252,878 g, 4°C, 3 h, then sampled using a Biocomp Gradient profiler (model 251) with constant monitoring of optical density at 254 nm using standard parameters. Data analysis was performed with Excel.

For sucrose gradient sedimentation analysis of nuclear lysates, thawed cells were suspended in 1 mL nuclear isolation buffer (10 mM HEPES-KOH pH 7.4, 100 mM KCl, 5 mM MgCl2 with 0.5% NP40, Roche Complete protease inhibitor cocktail tablets and Ribonuclease Inhibitors), and incubated on ice for 10 min, followed by centrifugation at 3,000 g, 4°C, 3 min. The nuclear pellets were suspended in 1 mL nuclear lysis buffer (20 mM HEPES-KOH pH 7.4, 300 mM KCl, 5 mM MgCl2, 1 mM DTT with 1% TritonX-100, Roche Complete protease inhibitor cocktail tablets and Ribonuclease Inhibitors), followed with sonication (50% intensity, 5 s pulse with 25 s rest, 5×, MICROSON XL 2000), and centrifugation at 21,130 g, 4°C, 10 min. The supernatants were normalized by A260 and layered onto 12 mL 10-50% sucrose gradients, centrifuged and sampled as described above.

### Crystal violet staining and cell cycle synchronization

4×10^4^ HEK 293T cells were seeded in 12-well plates in triplicate, and fixed in 10% formalin for 15 min at RT every 24 h. After washing with ddH_2_O, cells were stained with 0.1% crystal violet in methanol for 30 min at RT in the dark, followed with three ddH_2_O washes and dried. The cells were then immersed in 1 mL 10% acetic acid with shaking for 20 min. 20 µL of the solution was combined with 80 µL ddH_2_O in a 96-well plate, and the optical density at 590 nm was monitored with Synergy™ HT.

For cell cycle synchronization, 1.5×10^5^ HEK 293T cells were seeded in 12-well plates and cultured overnight. Cells were then treated with 2 mM thymidine for 16 h, followed with two PBS washes, and incubated with fresh media for 9 h before the second 2 mM thymidine block for 14 h following a previously published protocol^64^. Cells were washed with PBS, then incubated with fresh media to release from the G1/S boundary, and fixed in 10% formalin for immunostaining or harvested for western blotting.

### Northern blot

Total RNA was extracted from HEK 293T cells using TRIzol reagent. To determine changes in levels of LSU pre-rRNA intermediates, 3 µg of total RNA was run on a 1% agarose/1.25% formaldehyde gel in a 1.5 M tricine/ 1.5 M triethanolamine buffer, followed by an overnight transfer to a Hybond XL nylon membrane (GE Healthcare, RPN 303S) in 10× saline-sodium citrate transfer buffer after a brief 15 min soak in a 0.5 M sodium hydroxide solution. Membranes were then exposed to UV (254 nm) to immobilize the RNA, followed by incubation with denatured yeast tRNA for 1 h at 42°C, and hybridized overnight at 37°C with 5’ end radiolabeled oligonucleotide probe (P4 5’-CGGGAACTCGGCCCGAGCCGGCTCTCTCTTTCCCTCTCCG-3’) in a solution of 7.5× Denhardt’s solution, 5× sodium chloride-sodium phosphate-EDTA buffer with 0.1% SDS, as previously reported^59^. Membranes were also hybridized with a 7SL probe (7SL 5’-TGCTCCGTTTCCGACCTGGGCCGGTTCACCCCTCCTT-3’) as a loading control.

## Data availability

Proteomic data were deposited under accession PXD026880. During review, they can be accessed with username and password 7dkNTlia.

## Acknowledgements

We thank Franziska Bleichert and all members of the Slavoff and Baserga labs for helpful conversations. This work was supported by a Searle Scholars Program Award, an Odyssey Award from the Richard and Susan Smith Family Foundation, and start-up funds from Yale University West Campus (to S. A. S.). X.C. was supported in part by a Rudolph J. Anderson postdoctoral fellowship from Yale University. A.K. was in part supported by an NIH Predoctoral Training Grant (5T32GM06754 3-12). S.J.B, C.J.B. and C.M.H. were supported by R35 GM131687. C.M.H. was supported by an NSF GFRP.

## Author contributions

X.C., A.K., C.M.H., C.J.B., and S.Z. and designed and performed experiments and analyzed data. S. A. S. and S. J. B. designed experiments and analyzed data. X. C. and S. A. S. wrote the manuscript, and all authors edited and approved the final version of the manuscript.

