## Supplementary Information for "Nascent alt-protein chemoproteomics reveals a repressor of ribosome biogenesis"

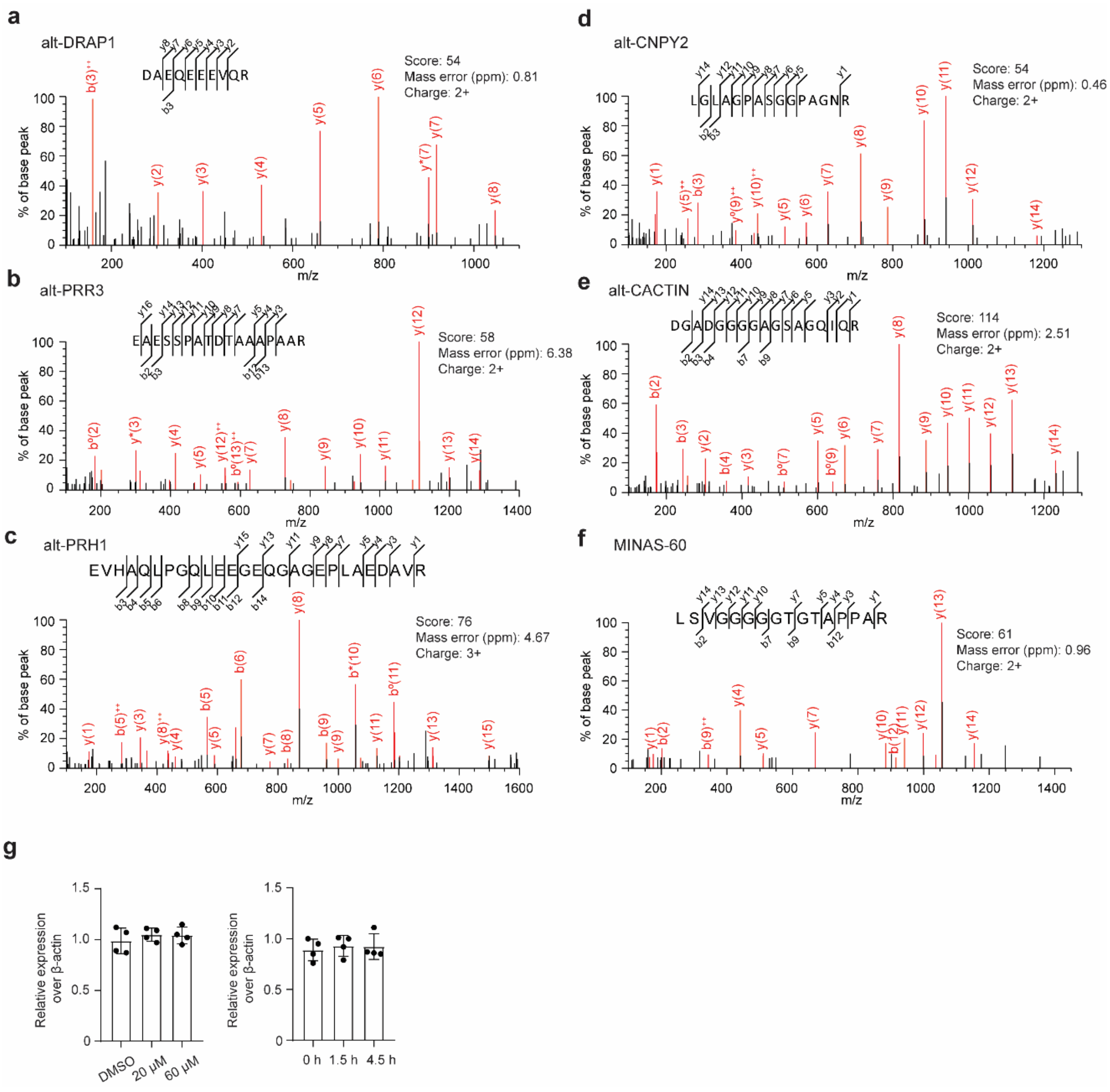


**Supplementary Figure 1. BONCAT-based chemoproteomic identification of newly synthesized alt-proteins. a-f** MS/MS spectrum of the tryptic peptide of alt-DRAP1 (**a**), alt-PRR3 (**b**), alt-PRH1 (**c**), alt-CNPY2 (**d**), alt-CACTIN (**e**), and MINAS-60 (**f**) detected by BONCAT-based chemoproteomic in HEK 293T cells. **g** Quantitative RT-PCR with primers specific to the *CNPY2* transcript variant 1 in HEK 293T cells treated with increasing amounts of etoposide or vehicle for 2 h (left), or with 60 μM etoposide or vehicle for different times (right). Error bars, standard error of the mean (s.e.m.), $N$ = 4 biologically independent samples, two-tailed $t$-test.

| | Location | Start codon | DMSO | Etoposide | Arsenite | Length (aa) | Evidences |
|---|---|---|---|---|---|---|---|
| alt-DRAP1 | 5'UTR | non-AUG | yes | | | 90 | None |
| alt-PRR3 | 5'UTR | AUG | yes | | | 93 | Proteomics and Ribo-seq |
| alt-PRH1 | 5'UTR | AUG | yes | | yes | 72 | Ribo-seq |
| alt-CNPY2 | 5'UTR | non-AUG | | yes | | 85 | None |
| alt-CACTIN | CDS | AUG | yes | | | 102 | Proteomics and Ribo-seq |
| MINAS-60 | CDS | AUG | yes | | | 130 | Predicted |

**Supplementary Table 1 | Summary of the six validated alt-proteins.** Evidences are provided by OpenProt_2021[1], alt-proteins detected by mass spectrometry are indicated with proteomics, and alt-proteins detected by ribosome sequencing are indicated with Ribo-seq.

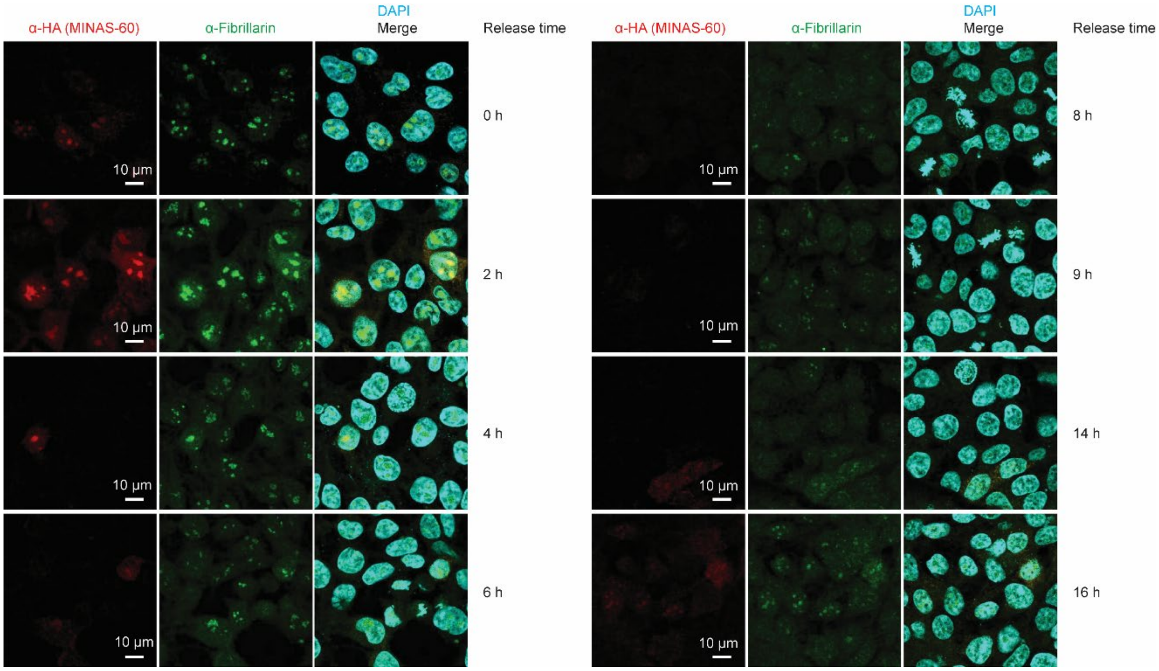


**Supplementary Figure 2. Nucleolar-localized MINAS-60 expression increases in early S phase.** Immunostaining analysis of synchronized MINAS-60 KI cells released from the G1/S boundary by a double thymidine block at the indicated time points with anti-HA (red), anti-fibrillarin (green), and DAPI (cyan). Scale bar, 10 μm. Data are representative of three biological replicates.

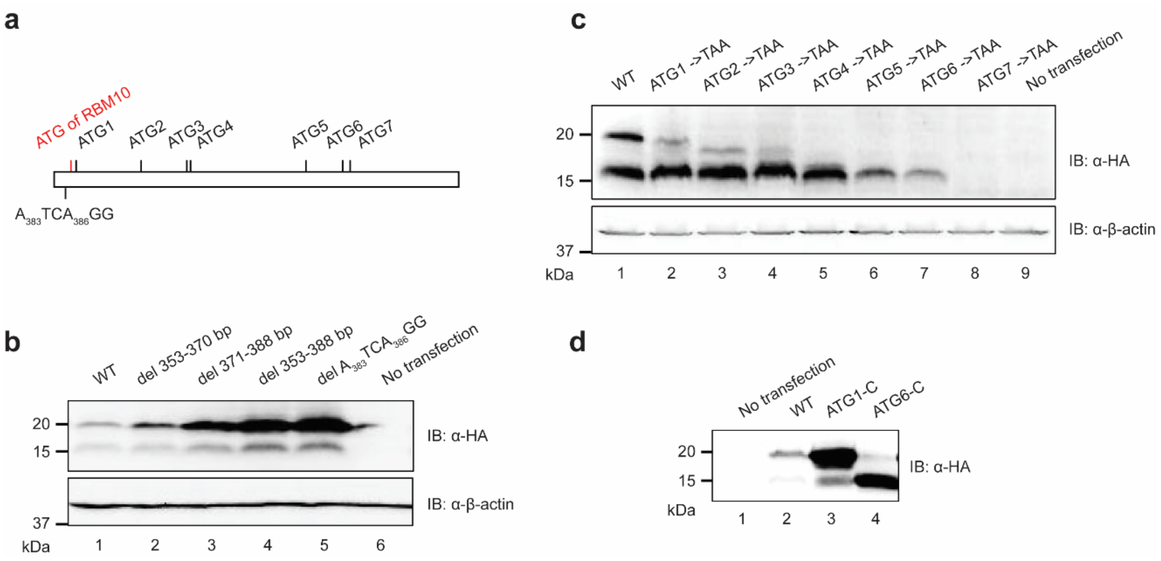


**Supplementary Figure 3. MINAS-60 is initiated from multiple start codons. a** Schematic representation of the region of human *RBM10* transcript variant 1 (tv 1) harboring candidate MINAS-60 start codons. The ATG start codon of RBM10 is indicated in red, and the two upstream non-ATG and seven internal ATG start codons that are upstream of and in-frame with the detected tryptic fragment of MINAS-60 are indicated in black. **b, c** Expression of a construct containing the full 5'UTR and wild-type MINAS-60 coding sequence (WT, lane 1), or MINAS-60 coding sequence with deletion (del, lanes 2-5, **b**), or MINAS-60 coding sequence with mutation (lane 2-8, **c**), indicated on the top derived from *RBM10* tv 1, with an HA tag appended to the C-terminus of MINAS-60, in HEK 293T cells was followed by western blotting with the antibodies indicated to the right. Untransfected (no transfection) HEK 293T cells served as a control. Data are representative of three biological replicates. **d** Expression of a construct containing the full 5'UTR and MINAS-60 coding sequence (lane 2), or ATG1 to the C-terminus of MINAS-60 (lane 3), or ATG6 to the C-terminus of MINAS-60 (lane 4) derived from *RBM10* tv 1, with an HA tag appended to the C-terminus of MINAS-60, in HEK 293T cells was followed by western blotting with the antibodies indicated to the right. Untransfected (no transfection) HEK 293T cells served as a control. Data are representative of three biological replicates.

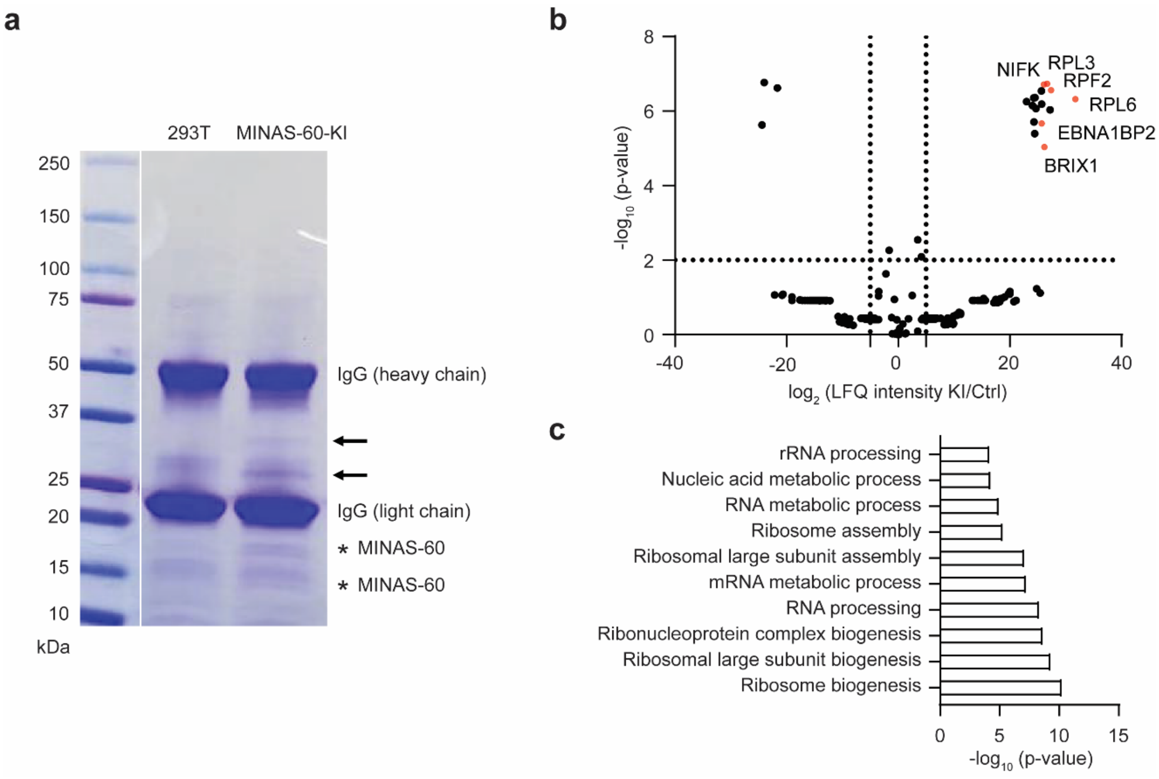


**Supplementary Figure 4. MINAS-60 associates with nucleolar LSU biogenesis factors. a** Coomassie blue staining of FLAG-IP from control (293T) or MINAS-60 KI (MINAS-60-KI) HEK 293T cells. Two major enriched bands from the KI FLAG-IP are indicated with black arrows, and MINAS-60 bands are indicated with stars. **b** Volcano plot of quantitative proteomics (*N* = 3) of gel bands (25-45 kDa) excised from MINAS-60 KI (KI) or control (Ctrl) HEK 293T nuclear lysate FLAG-IP samples after SDS-PAGE. LSU biogenesis factors are indicated in red and the gene names are labeled. For complete quantitative proteomics results, see Supplementary Data 3. **c** GO (biological processes) analysis of genes enriched (fold change ≥ 30) in MINAS-60 KI FLAG-IP over control with g:Profiler[2].

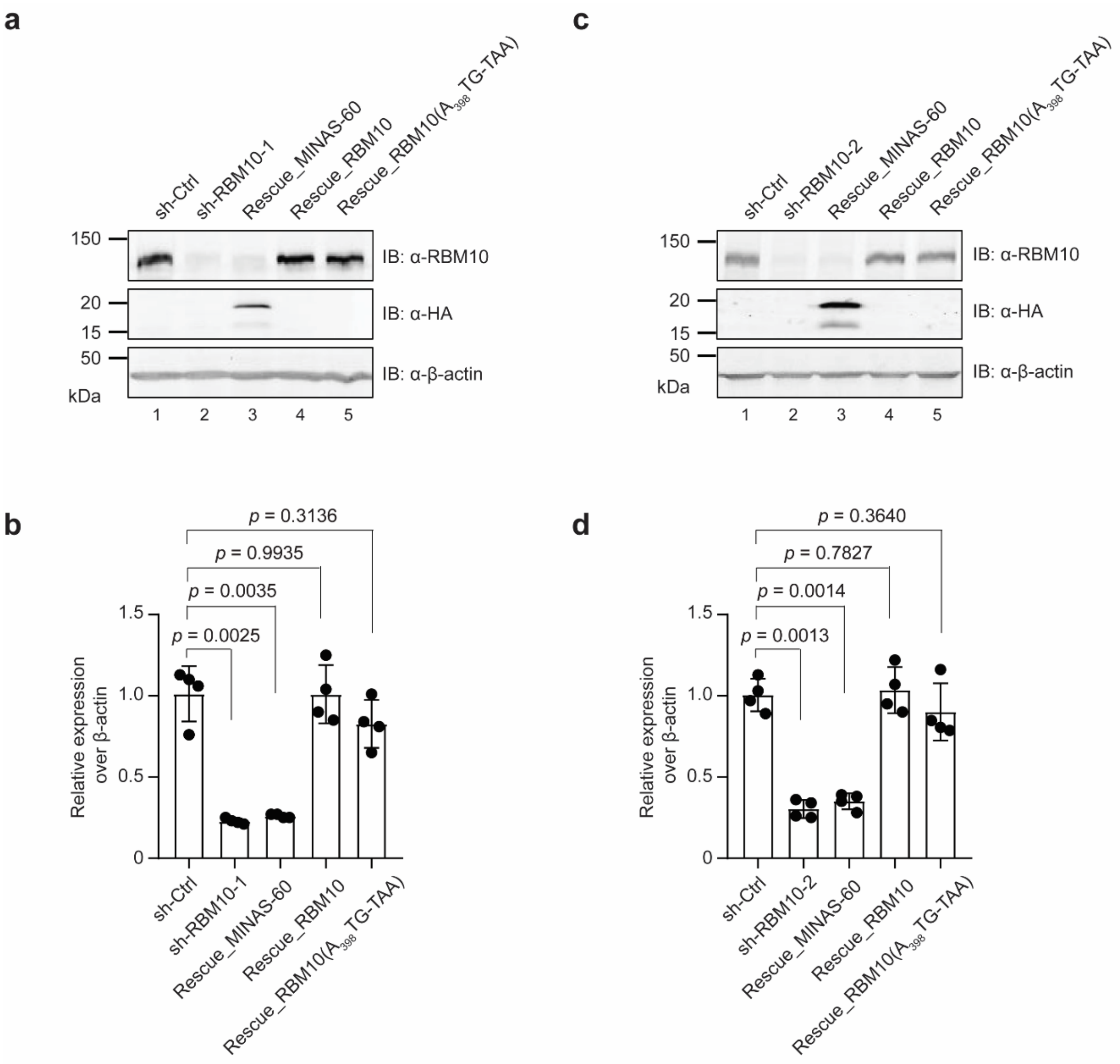


**Supplementary Figure 5. Validation of the *RBM10* knockdown and rescue HEK 293T cell lines. a, c** Western blot analysis of HEK 293T stably expressing empty pLKO.1 vector control (lane 1, sh-ctrl), one of the two *RBM10* shRNAs (lane 2, sh-RBM10-1 (**a**), sh-RBM10-2 (**c**)), rescue with MINAS-60 (lane 3, Rescue_MINAS-60), rescue with RBM10 (lane 4, Rescue_RBM10), or rescue with RBM10 bearing an $A_{398}$TG to TAA mutation (lane 5, Rescue_RBM10($A_{398}$TG-TAA)). Data are representative of three biological replicates. **b, d** Quantitative RT-PCR of the cell lines described above with primers specific to *RBM10* (error bars, standard error of the mean (s.e.m.)), $N = 4$ biologically independent samples, two-tailed *t*-test.

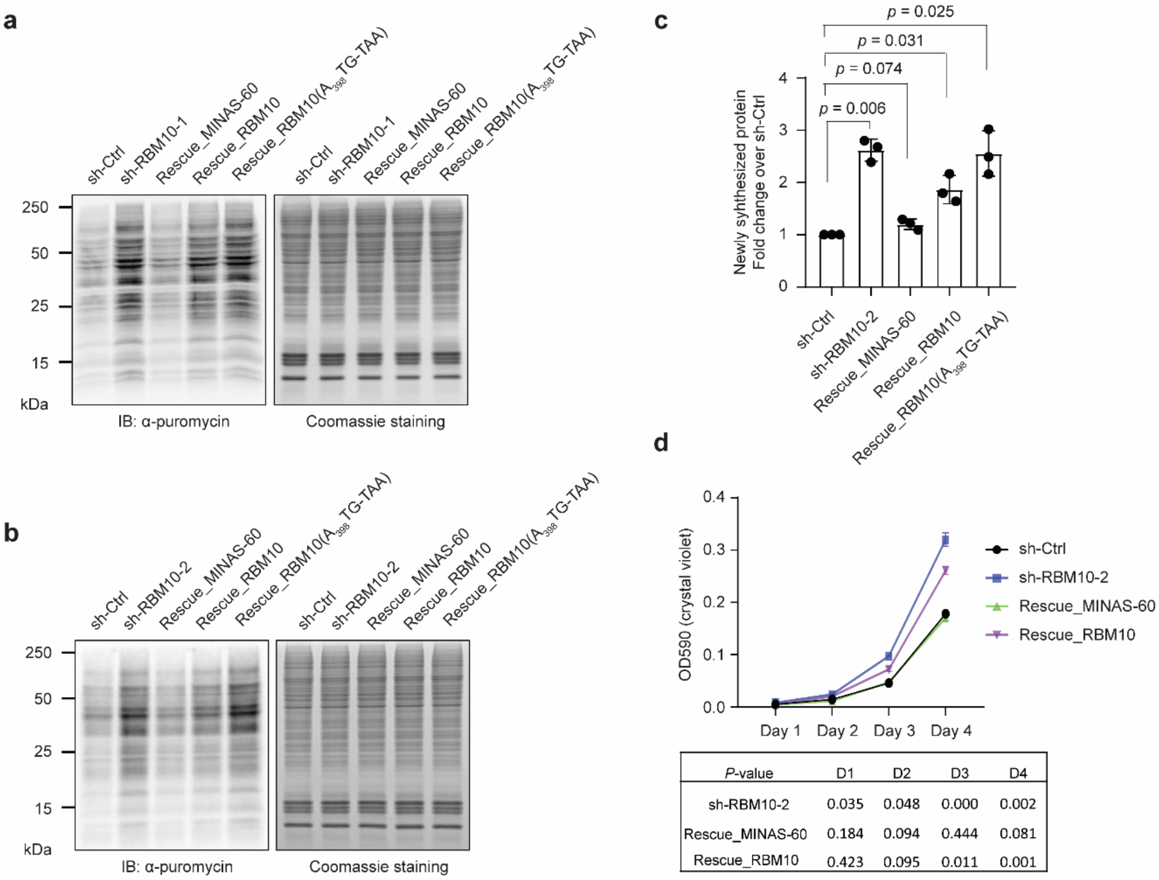


| *P*-value | D1 | D2 | D3 | D4 |
|---|---|---|---|---|
| sh-RBM10-2 | 0.035 | 0.048 | 0.000 | 0.002 |
| Rescue_MINAS-60 | 0.184 | 0.094 | 0.444 | 0.081 |
| Rescue_RBM10 | 0.423 | 0.095 | 0.011 | 0.001 |

**Supplementary Figure 6. MINAS-60 downregulates global protein synthesis and cell proliferation. a, b** HEK 293T cells stably expressing empty pLKO.1 vector control (sh-Ctrl), one of the two *RBM10* shRNA (sh-RBM10-1 (**a**) sh-RBM10-2 (**b**)), rescue with MINAS-60 (Rescue_MINAS-60), rescue with RBM10 (Rescue_RBM10), or rescue with RBM10 bearing an $A_{398}$TG to TAA mutation (Rescue_RBM10($A_{398}$TG-TAA)) were treated with 1 μM puromycin for 1 hour at 37℃ before harvesting and western blotting with anti-puromycin antibody. Coomassie staining served as a loading control. Data are representative of three biological replicates. **c** ImageJ was used to quantify the relative puromycin incorporation for cells indicated at the bottom relative to sh-Ctrl from three biological replicates. Data represent mean values ± s.e.m., and significance was evaluated with two-tailed *t*-test. **d** Growth curve of control (sh-Ctrl), *RBM10* knockdown with a second shRNA (sh-RBM10-2), rescue with MINAS-60 (Rescue_MINAS-60) and rescue with RBM10 (Rescue_RBM10) HEK 293T cells at the indicated number of days ($N = 3$). Data represent mean values ± s.e.m., and significance was evaluated with two-tailed *t*-test and shown below.

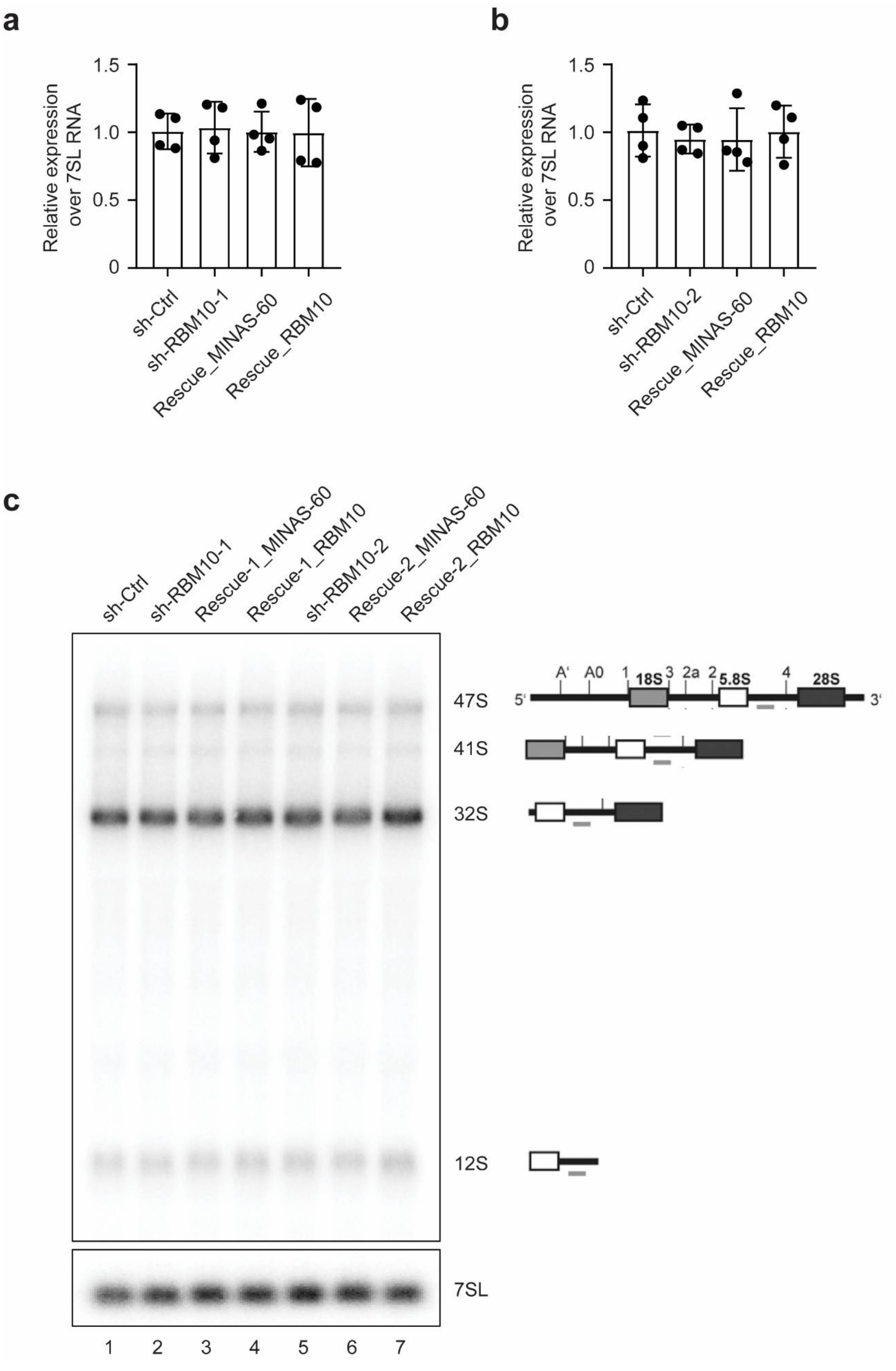
a
b
Relative expression over 7SL RNA
1.5
1.0
0.5
0
sh-Ctrl
sh-RBM10-1
Rescue_MINAS-60
Rescue_RBM10
sh-RBM10-2
c
sh-Ctrl
sh-RBM10-1
Rescue-1_MINAS-60
Rescue-1_RBM10
sh-RBM10-2
Rescue-2_MINAS-60
Rescue-2_RBM10
47S
41S
32S
12S
7SL
1 2 3 4 5 6 7
5'
3'
A'
A0
1
18S
3
2a
2
5.8S
4
28S

**Supplementary Figure 7. MINAS-60 does not regulate the transcription of pre-rRNA, nor the LSU pre-rRNA processing. a, b** Quantitative RT-PCR with primers specific to the primary pre-rRNA (47S/45S/30S) of HEK 293T stably expressing empty pLKO.1 vector control (sh-ctrl), one of the two *RBM10* shRNAs (sh-RBM10-1 (**a**), sh-RBM10-2 (**b**)), rescue with MINAS-60 (Rescue_MINAS-60), or rescue with RBM10 (Rescue_RBM10) (error bars, standard error of the mean (s.e.m.)), $N = 4$ biologically independent samples, two-tailed *t*-test. **c** Total RNA from HEK 293T stably expressing empty pLKO.1 vector control (lane 1, sh-ctrl), one of the two *RBM10* shRNAs (sh-RBM10-1 (lane 2), sh-RBM10-2 (lane 5)), rescue with MINAS-60 (Rescue-1_MINAS-60 (rescue on sh-RBM10-1 background, lane 3), Rescue-2_MINAS-60 (rescue on sh-RBM10-2 background, lane 6)), or rescue with RBM10 (Rescue-1_RBM10 (rescue on sh-RBM10-1 background, lane 4), Rescue-2_RBM10 (rescue on sh-RBM10-2 background, lane 7)) were isolated with TRIzol, and pre-rRNAs were separated by gel electrophoresis, followed with northern blotting using radioactively labeled P4 probe (gray lines, diagram at right). Northern blotting with a probe against the 7SL RNA was used as a loading control. Illustrations of the pre-rRNAs detected by P4 are indicated to the right of their respective bands. Data are representative of three biological replicates.

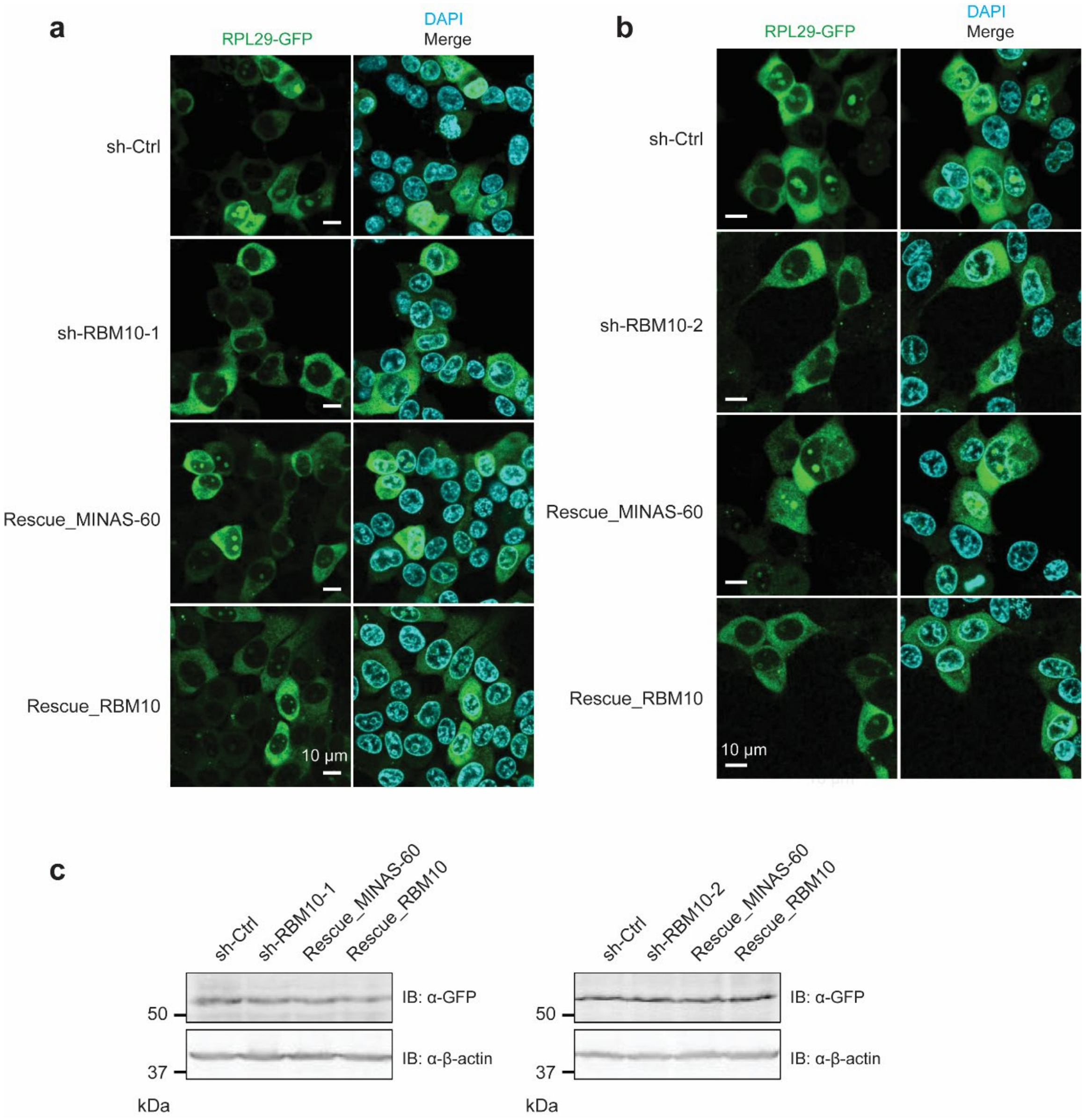


**Supplementary Figure 8. MINAS-60 downregulates LSU export. a, b** Confocal live-cell imaging of control (sh-Ctrl), *RBM10* knockdown with two independent shRNAs (sh-RBM10-1 (**a**), sh-RBM10-2 (**b**)), rescue with MINAS-60 (Rescue_MINAS-60) or rescue with RBM10 (Rescue_RBM10) HEK 293T cells stably expressing RPL29-GFP. Scale bar, 10 μm. Data are representative of three biological replicates. **c** Western blot of the cell lines described above with antibodies indicated on the right for comparison of RPL29-GFP expression. Data are representative of three biological replicates.

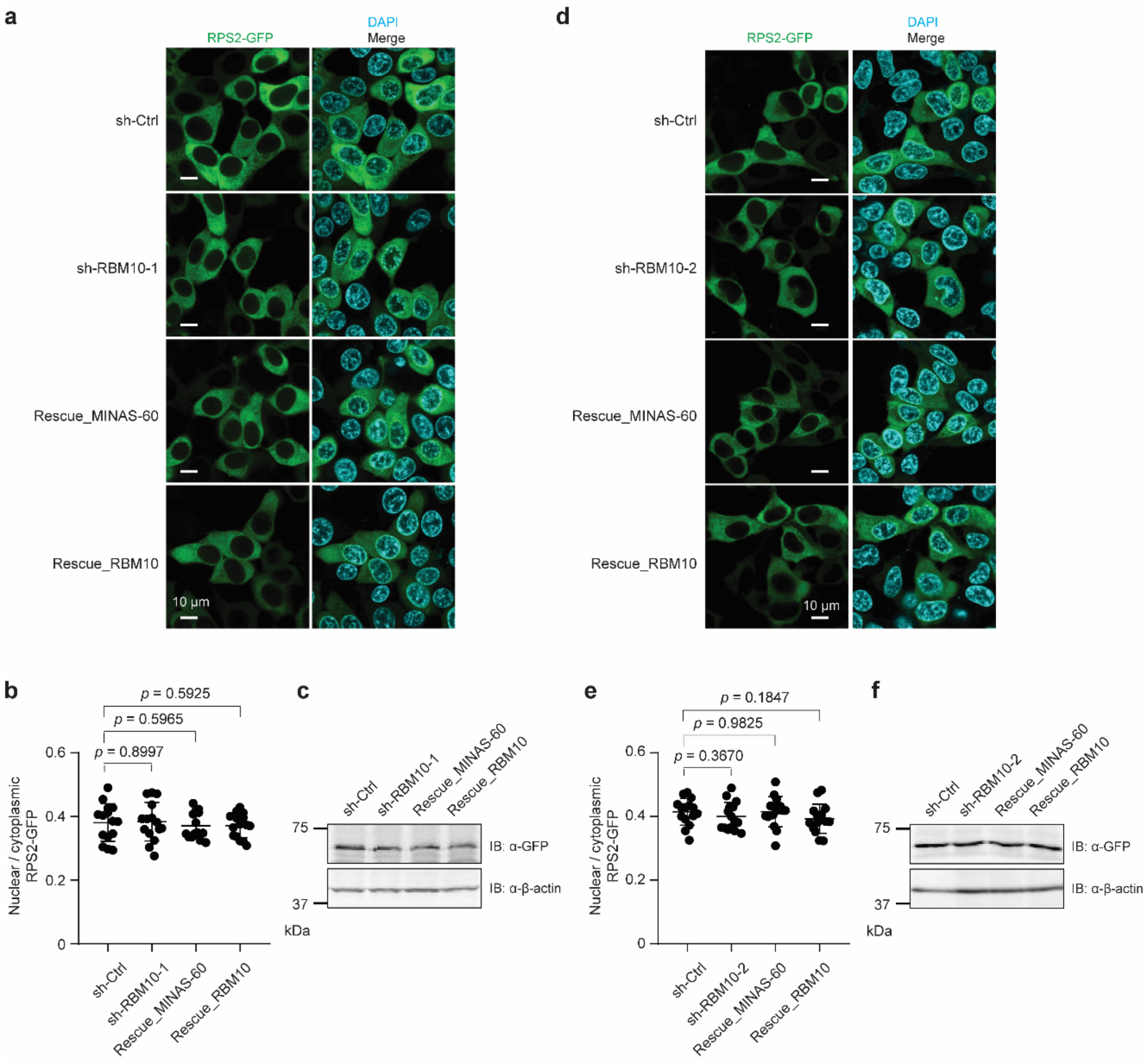


**Supplementary Figure 9. MINAS-60 does not regulate small 40S ribosomal subunit export. a, d** Confocal live-cell imaging of control (sh-Ctrl), *RBM10* knockdown with two independent shRNAs (sh-RBM10-1 (**a**) and sh-RBM10-2 (**d**)), rescue with MINAS-60 (Rescue_MINAS-60) or rescue with RBM10 (Rescue_RBM10) HEK 293T cells stably expressing RPS2-GFP. Scale bar, 10 µm. Data are representative of two biological replicates. **b, e** Quantitation of the RPS2-GFP signals in the cell lines described above. At least 13 fields of view were analyzed, totaling > 350 cells for each measurement. Data represent mean values ± s.e.m., and significance was evaluated with two-tailed *t*-test. **c, f** Western blot of the cell lines described above with antibodies indicated on the right for comparison of RPS2-GFP expression. Data are representative of two biological replicates.

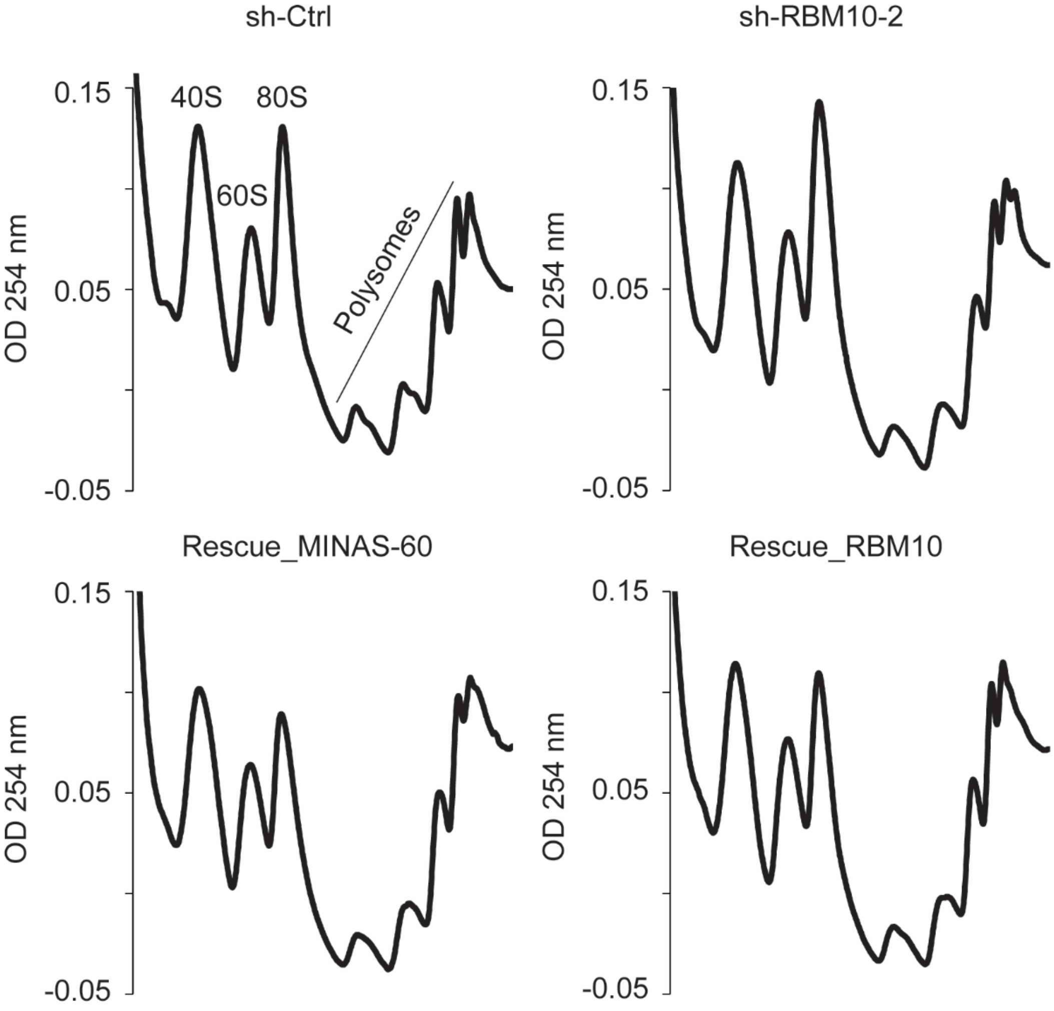


**Supplementary Figure 10. MINAS-60 downregulates LSU export.** Sucrose gradient sedimentation analysis of polysome fractions of cytoplasmic lysates from control (sh-Ctrl), *RBM10* knockdown with a second shRNA (sh-RBM10-2), rescue with MINAS-60 (Rescue_MINAS-60) or rescue with RBM10 (Rescue_RBM10) HEK 293T cells. Data are representative of three biological replicates.

| qRT-PCR primers (5' - 3') | | | |
|---|---|---|---|
| Target genes | Use | Forward primer | Reverse primer |
| CNPY2 | qRT-PCR | TTGATCCTTCCACCCATCGC | CATTGTCAGCCTCTCGGGAA |
| RBM10 | qRT-PCR | ATTTTGCGCAACCTGAACCC | GGTGGAGAGCTGGATGAAGG |
| Primary pre-rRNA | qRT-PCR | CTCCGTTATGGTAGCGCTGC | GCGGAACCCTCGCTTCTC |
| β-actin | qRT-PCR | AGGCACCAGGGCGTGAT | GCCCACATAGGAATCCTTCTGAC |
| 7SL | qRT-PCR | ATCGGGTGTCCGCACTAAGTT | CAGCACGGGAGTTTTGACCT |

**Supplementary Table 2 | qRT-PCR primers (5' - 3').**
